# Core Tri-fucosylation of Nematode N-glycans Requires Golgi α-mannosidase III Activity that Impacts Animal Growth and Behaviours

**DOI:** 10.1101/2024.07.18.600072

**Authors:** Jonatan Kendler, Florian Wöls, Saurabh Thapliyal, Elsa Arcalis, Hanna Gabriel, Sascha Kubitschek, Daniel Malzl, Maria R. Strobl, Dieter Palmberger, Thomas Luber, Carlo Unverzagt, Katharina Paschinger, Dominique A. Glauser, Iain B. H. Wilson, Shi Yan

## Abstract

Many nematodes possess N-glycans with complex core chitobiose modifications, which is a feature observed in various free-living and parasitic nematodes but is absent in mammals. Using *Caenorhabditis elegans* as a model to study N-glycan biosynthesis, we demonstrated that the core *N*-acetylglucosamine (GlcNAc) residues can be modified by three fucosyltransferases in the Golgi, namely FUT-1, FUT-6 and FUT-8. While the asparagine-linked GlcNAc is modified with a α1,3- and α1,6-linked fucose by FUT-1 and FUT-8 respectively, the distal GlcNAc residue is α1,3-fucosylated solely by FUT-6. Interestingly, FUT-6 can only fucosylate N-glycan structures lacking the α1,6-mannose upper arm, indicating that a specific α-mannosidase is required to generate substrates for subsequent FUT-6 activity. By analysing the N-glycomes of *aman-3* mutants (tm5400 and a CRISPR/Cas9 knockout, *hex-2;hex-3;aman-3*) using offline HPLC-MALDI-TOF MS/MS, we observed that the absence of the *aman-3* gene abolishes α1,3-fucosylation of the distal GlcNAc of N-glycans, which suggests that AMAN-3 is the relevant mannosidase on whose action FUT-6 depends. To further investigate it, we recombinantly expressed AMAN-3 in insect cells and characterised its enzymatic activity *in vitro*. In contrast to the classical Golgi α-mannosidase II (AMAN-2), AMAN-3 displayed a cobalt-dependent α1,6-mannosidase activity towards N-glycans. Using AMAN-3 and other recombinant *C. elegans* glycoenzymes, we remodelled a fluorescein conjugated-Man_5_GlcNAc_2_ structure; we were able to mimic N-glycan biosynthesis in the Golgi and generate a tri-fucosylated glycan *in vitro*. We performed confocal microscopy studies using a knock-in strain (*aman-3*::*eGFP*) and could show the Golgi localisation of AMAN-3. In addition, using a high-content computer-assisted *C. elegans* analysis platform, we observed that AMAN-3 deficient worms display significant developmental delays, morphological and behavioural alterations in comparison to the wild type. Therefore, our data suggested that AMAN-3 participates in nematode N-glycan biosynthesis in the Golgi and generates substrates for FUT-6; thereby, this enzyme is essential for the formation of the unusual tri-fucosylated chitobiose cores of nematode N-glycans, which may play important roles in nematode development and behaviour.

**Background:** Tri-fucosylation of N-glycan core is a conserved feature seen in the N-glycomes of several nematode species. However, beyond the three core fucosyltransferases, we know very little about the biosynthesis and biological function of these core modifications.

**Results:** Comparative glycomics data revealed that *aman-3* mutants possess underfucosylated N-glycomes. Biochemical characterisation of AMAN-3 clarified its optimal reaction conditions and substrate specificity and, we demonstrated a Golgi localisation. Thereafter in vitro reconstruction of biosynthesis of a core tri-fucosylated N-glycan was achieved using 8 recombinant *C. elegans* glycoenzymes. Notably, *aman-3* deficient worms exhibited significant developmental and behavioural changes.

**Conclusion:** AMAN-3 is a Golgi α-mannosidase required for core fucosylation of the distal *N*-acetylglucosamine of N-glycoproteins.

**Significance:** This study elucidates the key role of a novel Golgi α-mannosidase in the biosynthesis of the unusual N-glycans of *C. elegans* and related nematodes, thereby setting the stage for new approaches to study the roles of glycan in the biology and immunology of nematode glycoproteins.

## Introduction

Protein glycosylation is a ubiquitous and evolutionarily conserved post-translational modification observed across diverse species. It exerts pleiotropic effects on many biological phenomena under both physiological and pathophysiological conditions. As an important modulator of signalling, cell adhesion and cell-cell interactions, protein glycosylation profoundly influences embryogenesis, tissue homeostasis, as well as cancer progression [1–3]. The nervous system also relies on precise glycosylation for proper neuronal development and function [4]. Moreover, glycoconjugates of pathogens are the major determinants that trigger immune responses, as they often possess ligands for innate immune recognition and promote production of anti-glycan antibodies [5, 6].

*Caenorhabditis elegans* has been employed as a good model to study the glycosylation patterns conserved among parasitic species. In contrast to mammals, it has been shown that *C. elegans* can express complicated core modified N-glycan structures and a portion of these structural elements (glyco-epitopes) can also be found in parasitic nematodes [7]. Some nematode glyco-epitopes are obviously immunogenic in mammals, for instance the anti-HRP epitope (core α1,3-fucose) is recognised by IgE and IgG antibodies of the host [8, 9].

Correct glycosylation of an antigen is considered, in addition to the protein backbone, an important factor for the antigenicity of parasite proteins. Therefore, identifying the correct glycoepitopes and glycan structures on the native glycoproteins and characterising the glyco-enzymes involved in the biosynthesis are two major challenges for glycobiologists.

In our previous work, we have proven the function of the three fucosyltransferases (FUTs), which are involved in the biosynthesis of the highly fucosylated N-glycan core in *C. elegans*: FUT-1 and FUT-8 respectively direct the α1,3- and α1,6-fucosylation of the proximal GlcNAc residue [10, 11], whereas FUT-6 is responsible for α1,3-fucosylation of the distal GlcNAc residue (summarised in ***Figure 1***) [12]. Homologues of *C. elegans* FUT-8 have been well studied in both vertebrates and invertebrates [13, 11, 14]; enzymes with the same function as FUT-1 are characterised from plants and invertebrates [15], whereas FUT-6 [12] is a unique core fucosyltransferase occurring in some nematodes. It is noteworthy that FUT-6 displayed a strong bias against the presence of the α1,6-mannose residue (known as the ‘upper-arm’) in contrast to the α1,3-mannose residue (known as the ‘lower-arm’) of N-glycan structures as judged by the results of *in vitro* enzymatic assays [12]; in other words, FUT-6 only fucosylated the structures lacking the α1,6-linked mannose. Analysing the glycan structures from wild type and *fut-6* deficient mutants as well as a double hexosaminidase mutant (*hex-2;hex-3*) yielded further evidence for this property of FUT-6 [16].

**Figure 1.**
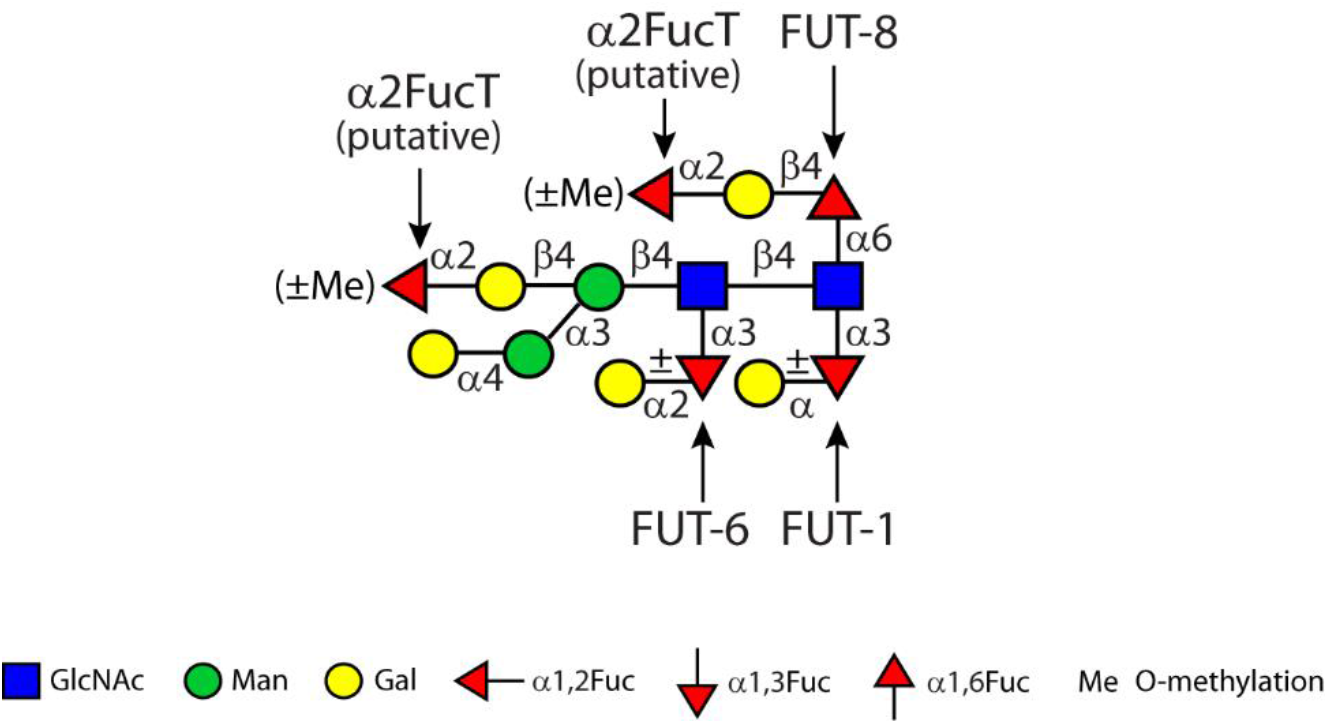
An illustration of fucose-rich glycans of the wild type *C. elegans* using the Symbol Nomenclature for Glycans (SNFG) [48]. Fucosyltransferases (FUTs) responsible for the modification of core fucose with a specific glycosidic linkage are indicated with arrows.

In addition to the core difucosylation of the proximal GlcNAc residue, distal GlcNAc fucosylation seems to be conserved in a number of nematode species, such as *Pristionchus pacificus, Ascaris suum, Oesophagostomum dentatum* and *Haemonchus contortus* [17, 16, 18, 19], probably due to the activity of FUT-6 orthologues, but are absent from filarial worms or *Trichuris suis* [20, 9]. In the case of *Haemonchus*, a predicted glycosyltransferase (Genbank® accession number: CDJ84058.1) possesses the highest homology to *C. elegans* FUT-6 in comparison to other FUT homologues. Consistent with the *in vitro* properties of FUT-6 (see above), detailed N-glycan structural analyses of these nematodes suggested the absence of the α1,6-mannose residue on the glycans which carry either solely an α1,3-linked fucose or a galactosylated fucose disaccharide unit on the distal GlcNAc. Presumably, the ‘upper arm’ generates a steric hindrance which restricts the access of FUT-6 to the 3-OH position of the distal GlcNAc. These observations led to the assumption that a Golgi α1,6-mannosidase activity must be essential for a proper biosynthesis of such structures prior to further processing by FUT-6 homologues.

There are three glycosyl hydrolase GH38 homologues in *C. elegans*: one lysosomal (AMAN-1), one classical Golgi mannosidase II (AMAN-2) and a hypothetical mannosidase (AMAN-3). In insects, enzymes with similarities to the latter (mannosidase III or ManIIb) have been characterised and shown to have Co(II)-dependent activities towards N-glycans [21, 22]. In contrast to AMAN-2 that unequivocally impacts *C. elegans* N-glycan biosynthesis, it remained unclear if AMAN-3 can process N-glycoproteins despite its activity towards an artificial substrate in the presence of Cobalt(II) chloride [23]. By BLAST search using the extracellular domain of *C. elegans* AMAN-3, α-mannosidase III homologues can also be found in *H. contortus* (CDJ83252.1) and *A. suum* (ERG79326.1) with 39% and 44% identity respectively. Therefore, it is highly possible that in addition to FUT-1 and FUT-8 homologues, these nematodes also express α-mannosidase III and FUT-6 homologues, which may sequentially act on N-glycans to create the highly fucosylated core structures found in these species.

In this study, we systematically investigated the enzymatic properties of the *C. elegans* AMAN-3 and proved its involvement in N-glycan biosynthesis. AMAN-3 deficiency not only impacts the core fucosylation patterns of nematode N-glycoproteins, but also alters animal development and results in behavioural changes.

## Results

### α1,6-specific mannosidase activity is lacking in aman-3 mutant worm lysates

In a previous study, we showed that the unpurified recombinant product of *C. elegans* F48C1.1 or *aman-3* gene could degrade *p*-nitrophenyl-α-mannoside and potentially cause degradation of pyridylaminated Man_5_GlcNAc_2_ [23]. In order to confirm the Co(II)-dependent mannosidase activity of native AMAN-3, verify its sensitivity to the swainsonine inhibitor and better delineate its N-glycan substrate specificity, we conducted additional analyses to assess how a selected N-glycan (Man_3_GlcNAc_2_, PA-MM in ***Figure 2***) is processed by lysates from wild-type worms (N2), *aman-3* loss-of-function mutants (tm5400), and *hex-2;hex-3* mutants with deletions in two Golgi hexosaminidase genes [16]. Noteworthily, the *hex-2;hex-3* mutant is known to possess high amount of N-glycan structures missing the α1,6-mannose upper-arm, presumably due to a high α1,6-specific mannosidase activity in this mutant. At approximately neutral pH, a removal of one mannose from PA-MM was observed with N2 and *hex-2;hex-3* lysates (***Figure 2D*** and ***F***), but not for the *aman-3* lysate (***Figure 2H-I***); this is evidenced by the appearance of a new HPLC peak at 5.7 g.u. and confirmed by MALDI-TOF-MS. Considering the forward shift in elution time, the Man_2_GlcNAc_2_ product was concluded to be Manα1,3Manβ1,4GlcNAcβ1,4GlcNAc-PA [24]. We conclude that the *aman-3* mutant lacked an Co(II)-dependent, swainsonine-inhibitable α1,6-specific mannosidase activity.

**Figure 2.**
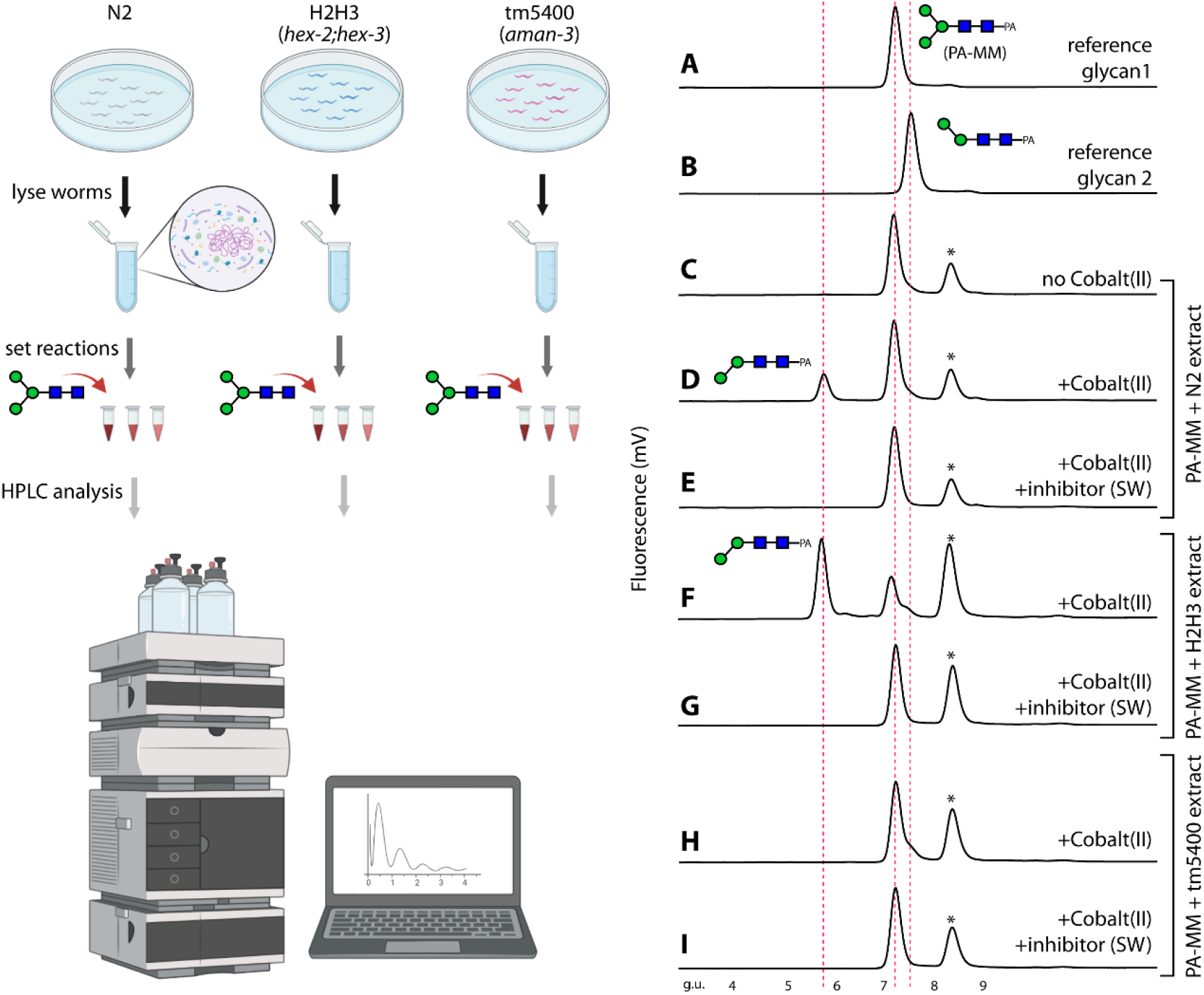
Detection of AMAN-3 activity in the crude extracts of different *C. elegans* strains. Worms were lysed to release native enzymes and clear supernatants were used to digest a pyridylamino-labelled N-glycan under various conditions; reaction mixtures were analysed on reversed phase HPLC (left panel, created with BioRender.com). A selected N-glycan (**A**, PA-MM, 7.2 g.u.) was incubated with worm supernatants prepared from either the wild type (N2; **C-E**), *hex-2;hex-3* double knockout (H2H3; **F** and **G**) or an *aman-3* single knockout (tm5400; **H** and **I**) with/without the presence of Cobalt (II) or Cobalt (II) plus swainsonine (SW). The presence of an early-eluting product, missing the α1,6-linked mannose, on HPLC chromatograms (**D** and **F**) indicated the α-mannosidase III activities. This Cobalt (II)-dependent demannosylation was only observed when N2 or H2H3 extract was used but was absent when the tm5400 extract was used (**H**). Peaks marked with an asterisk contain non-glycan contaminants; g.u. is an abbreviation of glucose units. The peaks were all analysed by MALDI-TOF MS and the major Co(II)-dependent product at 5.7 g.u. shown to have a glycan of *m/z* 827 [M+H]^+^ as compared to the substrate of *m/z* 989.

### N-Glycomes of aman-3 deficient mutants

Many N-glycans in wild-type *C. elegans* lack the α1,6-mannose residue as well as a ‘lower arm’ β1,2-GlcNAc residue on the α1,3-mannose [25–27], whereas the *hex-2;hex-3* double mutant has high amounts of glycans lacking the α1,6-mannose, but presenting the non-reducing GlcNAc residue [16]; on the other hand, the *bre-1* mutant lacks fucosylated glycans due to a mutation in the GDP-Man dehydratase gene [28]. Therefore, considering the hypothesis that AMAN-3 was an α1,6-specific mannosidase, we compared wild-type (N2), *hex-2;hex-3* and *bre-1* strains with an *aman-3* mutant (tm5400) and a *hex-2;hex-3;aman-3* triple mutant (cop1842). Distinct and major shifts in the PNGase A-released N-glycome were observed for all mutants examined and up to three fucose residues were detected in the overall MALDI-TOF MS profile for the *aman-3* mutant but only one fucose in the *hex-2;hex-3;aman-3* triple mutant (***Figure 3***), whereby the major N-glycans in these two strains were respectively Hex_5_HexNAc_2_Fuc_2-3_ and Hex_4_HexNAc_3_Fuc_1_.

**Figure 3.**
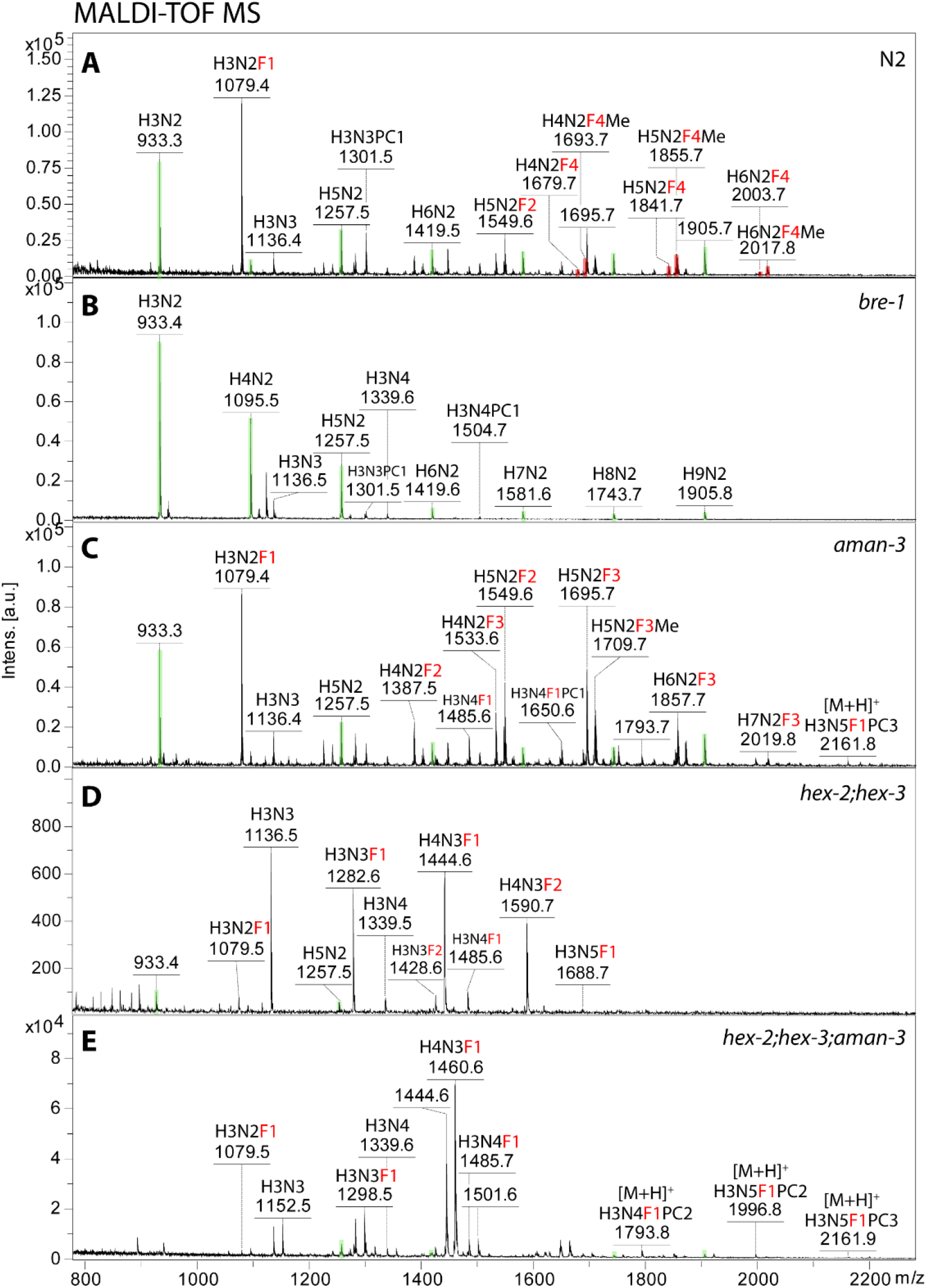
MALDI-TOF MS spectra of native N-glycans from the wild type and knockout strains. N-glycans released by PNGase A were subject to solid phase extraction and MS analysis. Most of the glycans present on the spectra are in [M+Na]^+^ and [M+K]^+^ forms except for PC-modified glycans detected partly in [M+H]^+^ forms. Peaks are annotated with *m/z* values and glycan compositions (H, hexose; N, *N*-acetyl-hexosamine; F, fucose, highlighted in red; Me, methyl group; PC, phosphorylcholine). Peaks indicating pauci- and oligo-mannosidic glycans are highlighted in green. In comparison to the N2 wild type that possesses tetrafucosylated N-glycans (**A**), underfucosylation with maximal 3 fucoses is observed in *aman-3* single mutant (**C**). In comparison to the H2H3 double mutant (**D**) [16], *hex-2;hex-3;aman-3* triple knockout (**E**) possesses three major compositions (H_3-4_N_3_F_0-1_) and the difucosylated glycan H_4_N_3_F_2_ (*m/z* 1590) is absent on the spectrum (the 4^th^ and 5^th^ panels). *Bre-1* [28] is a fucose-free mutant deficient in GDP-mannose 4,6-dehydratase (**B**).

To investigate the N-glycomic changes more exactly, HPLC was performed on a fused core RP-amide column and fractions were individually subject to MALDI-TOF MS/MS (***Figure 4***). The major glycan in the triple mutant (Hex_4_HexNAc_3_Fuc_1_; *m/z* 1500 as a pyridylaminated glycan) possessed a ‘GalFuc’ motif, due to β1,4-galactosylation of the core α1,6-fucose by GALT-1 [29], resulting in a strong Y1 fragment at *m/z* 608 (***Figure 4b***). The traces of glycans with two fucose residues were due to α1,2-fucosylation of the GalFuc (Y1 fragment at *m/z* 754), an epitope previously found in some fucosyltransferase mutants [30]. The high degree of substitution of the α1,3-mannose by the β1,2-GlcNAc residue in the triple mutant is probably the reason for the lack of methylation or α-galactosylation of the α1,3-mannose as well as the absence of bisecting galactose or reducing terminal core α1,3-fucose found in wild-type N-glycomes [31, 25]; in contrast, methylation of the α1,6-mannose was observed (see glycans of *m/z* 1206 and 1514; ***Figure 4***). As complex glycosylation was not blocked, but perhaps even preferred, a range of phosphorylcholine-modified N-glycans were also detected in the triple mutant (***Figure 4***). Only residual amounts of glycans lacking the α1,6-mannose (late-eluting GalFuc-modified isomers of *m/z* 1135 and 1338) were observed.

**Figure 4.**
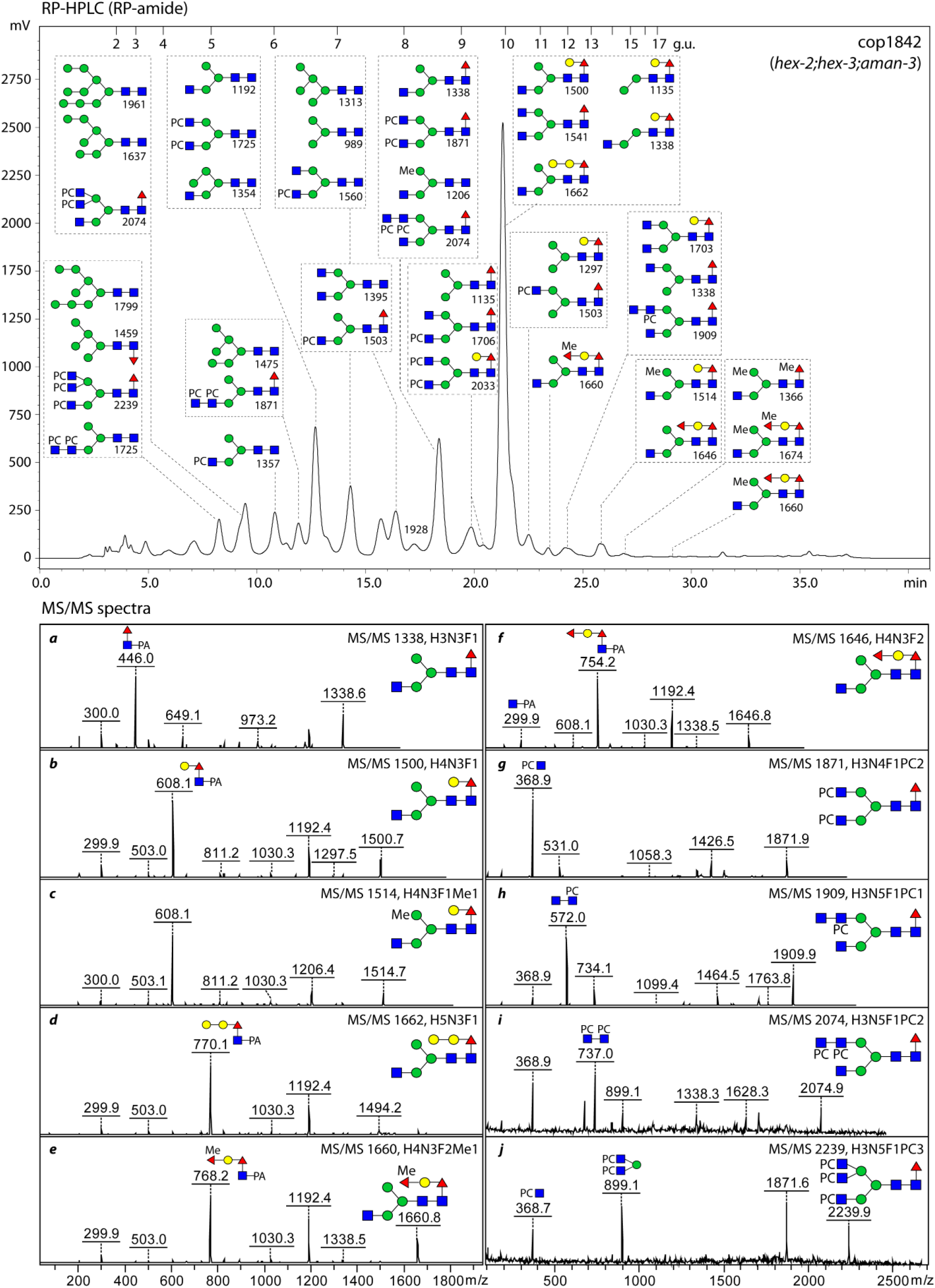
RP-HPLC chromatogram and MS/MS spectra of PNGase A-released N-glycans from the *hex-2;hex-3;aman-3* triple knockout (cop1842). PA-labelled glycans were fractionated on an RP-amide column and analysed by MALDI TOF MS/MS. Based on their elution patterns (glucose units) and MS data, predominant glycan structures detected in major HPLC peaks and key B/Y fragment ions are annotated with SNFG format and *m/z* values ([M+H]^+^). Structural assignments are on the basis of MS/MS as well as comparisons to other studies using the same HPLC column [19, 27].

A comparison of HPLC chromatograms of N2 and *aman-3* single mutant demonstrated shifts of major peaks (***Supplementary Figure 3***). Tri- and tetra-fucosylated structures were observed between 10 and 14 minutes (5.5-7.0 g.u.). While tetra-fucosylated glycans were observed in N2 (Hex_4-6_HexNAc_2_Fuc_4_Me_0-1_), these were completely absent in tm5400 and tri-fucosylated structures (Hex_4-7_HexNAc_2_Fuc_3_Me_0-1_) were more pronounced in this strain (peak ***a, b*** and ***i*** to ***v*** in ***Supplementary Figure 3***). In comparison to N2, two peaks containing primarily Man_3_GlcNAc_2_, Man_5_GlcNAc_2_ and Man_3_GlcNAc_2_Fuc_1_ structures were decreased in tm5400 (peak ***c*** and ***d*** in ***Supplementary Figure 3***).

### In vitro enzymatic activity and intracellular localisation of AMAN-3

The impact of deleting *aman-3* on the N-glycome, especially in the *hex-2;hex-3* background, directed us to more specifically examine the biochemical characteristics of the recombinant AMAN-3, which was expressed in Sf9 cells. His-tagged recombinant AMAN-3 was purified and its sequence was verified by LC-MS/MS (**Supplementary Table 1**). The Co(II)-dependence and slightly acidic pH optimum (6.5) were confirmed (***Figure 5A*** and ***B***); AMAN-3 retained over 70% activity in a broad temperature range between 10°C and 30°C, whereas the activity quickly declined under temperatures above 30°C (***Figure 5C***). The pH optimum data as well as glycomic data strongly suggest that AMAN-3 is a Golgi-resident enzyme. To verify this, confocal microscopy was employed to examine the distribution of GFP-fused AMAN-3 in live *aman-3::egfp* worms. Despite the low expression of AMAN-3, micrographs indicated that AMAN-3 signals tend to overlap with fluorescent signals of BODIPY TR ceramide, a dye used to stain Golgi apparatus (***Figure 5D***).

**Figure 5.**
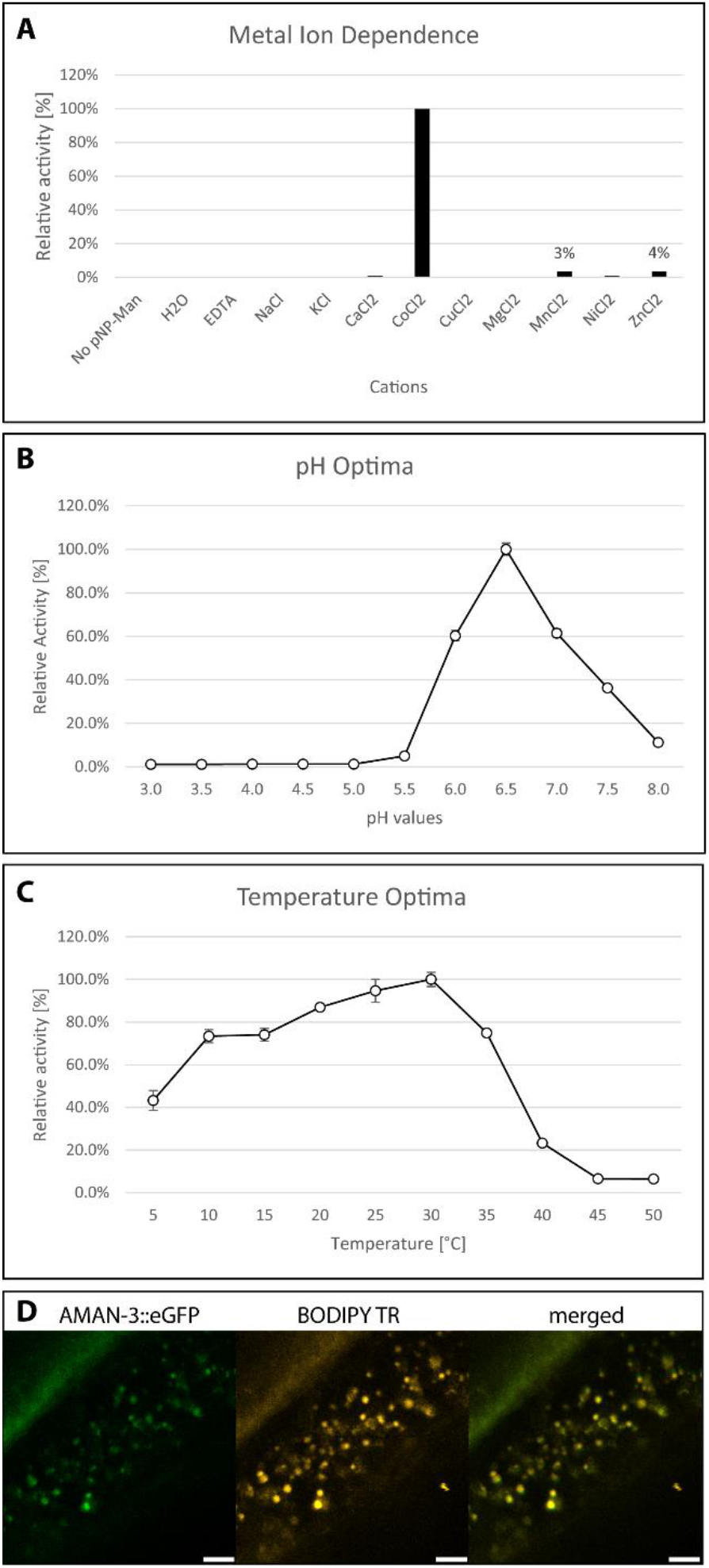
Enzymatic characterisation of recombinant AMAN-3 and confocal micrographs of AMAN-3::eGFP worm. Analyses were performed in 96-well microtiter plates by incubating purified AMAN-3 with pNP-α-Man under various conditions. OD405 absorption of the reaction mixtures was measured, and relative activities were calculated by comparing obtained values with the highest value in each experiment. Impact of different metal cations on AMAN-3 was investigated, which indicated that CoCl_2_ is a strong activator of AMAN-3 (**A**). The pH optimum of AMAN-3 is at 6.5 (**B**) and temperature optimum is at 30°C (**C**). Values represent averages ± standard errors (*n* = 4). (**D**) Confocal images of a paralysed adult worm demonstrate the overlapped AMAN-3::eGFP signal (green) and BODIPY signal (used to stain Golgi apparatus, shown in gold). White bars at the bottom indicate the length of 5 micrometres.

In terms of substrate specificity, we assayed the classical Golgi AMAN-2 and novel AMAN-3 using HPLC purified N-glycans with defined structures. Only AMAN-3 removed one mannose residue from Man_3_GlcNAc_2_, Man_5_GlcNAc_2_, Man_3_GlcNAc_3_, and Man_5_GlcNAc_3_ as indicated by a mass shift of 162, whereas AMAN-2 only digested Man_5_GlcNAc_3_, removing two mannose residues (***Figure 6*** *and* ***Supplementary Figure 4***). To further investigate which substrate is favoured by AMAN-3, we set reactions at room temperature with equal amount of substrate and quantified the products on HPLC. Data indicated that AMAN-3 removed solely the α1,6-mannose residue from these glycan substrates and except for Man_3_GlcNAc_3_ resulted in shifts in HPLC retention time (***Figure 6***). Full conversion was observed on Man_3_GlcNAc_2_, whereas for Man_5_GlcNAc_2_, Man_3_GlcNAc_3_ and Man_3_GlcNAc_3_ partial conversions (74.0%, 50.0% and 36.5%, respectively) were observed (***Figure 7***). The partial demannosylation from Man_3_GlcNAc_3_ to Man_2_GlcNAc_3_ by AMAN-3 was verified by MALDI-TOF MS and MS/MS (***Supplementary Figure 5***).

**Figure 6.**
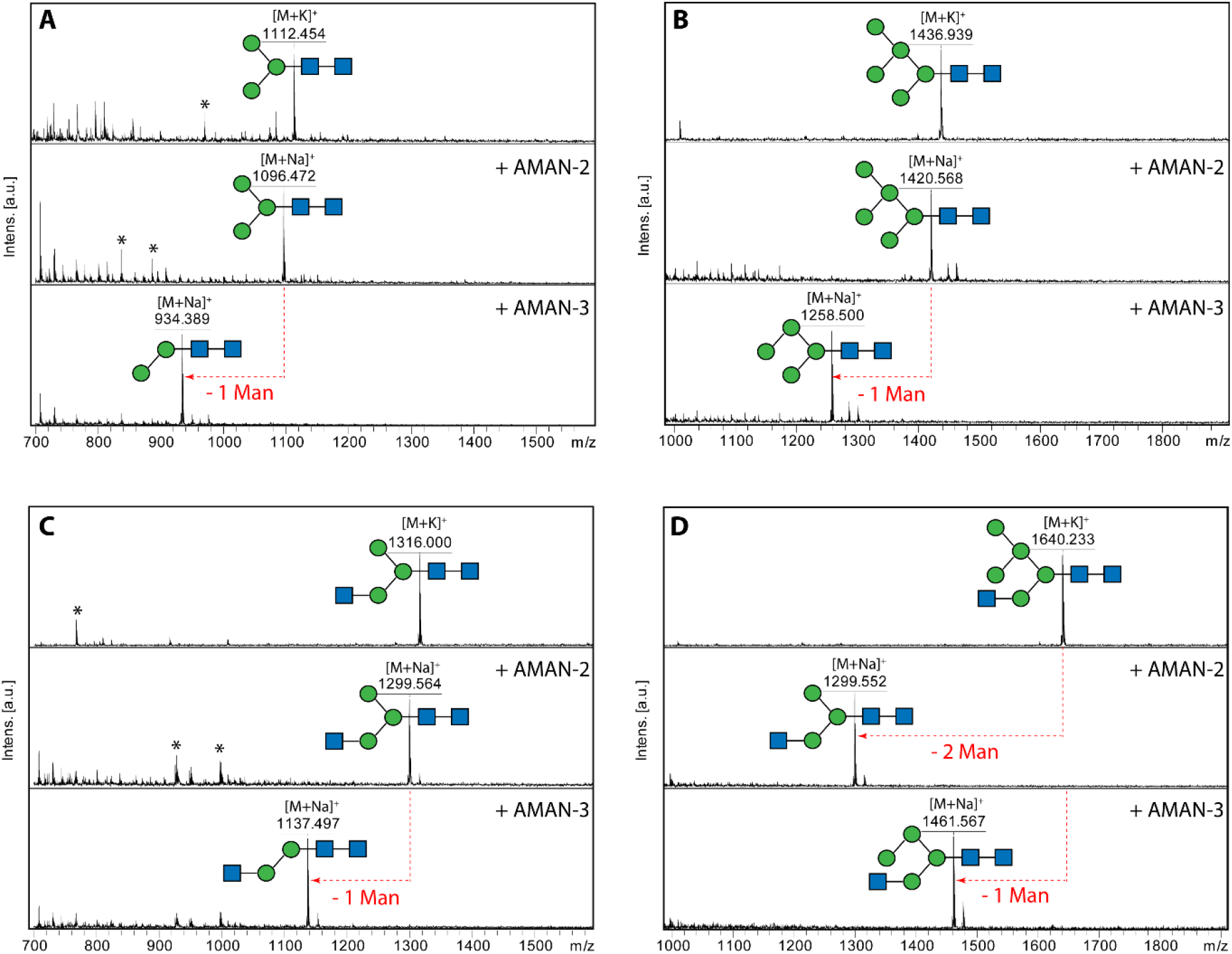
A comparison of substrate specificities between AMAN-2 and AMAN-3. Four RP-HPLC purified AEAB-labelled N-glycans (Man_3-5_GlcNAc_2-3_) were incubated with the recombinant mannosidases for 24 hours under their optimal conditions. Reaction mixtures were measured by MALDI-TOF MS and MS/MS. The two paucimannosidic structures Man_3_GlcNAc_2_ (**A**, *m/z* 1112) and Man_5_GlcNAc_2_ (**B**, *m/z* 1436) as well as a truncated complex structure Man_3_GlcNAc_3_ (**C**, *m/z* 1316) are substrates of AMAN-3 but not AMAN-2, as indicated by the loss of solely the α1,6-linked mannose residue. AMAN-2 removes two mannose residues from the hybrid N-glycan Man_5_GlcNAc_3_ (**D**, *m/z* 1640), whereas AMAN-3 selectively removes the terminal α1,6-linked mannose from it. Peaks marked with asterisks are non-glycan impurities.

**Figure 7.**
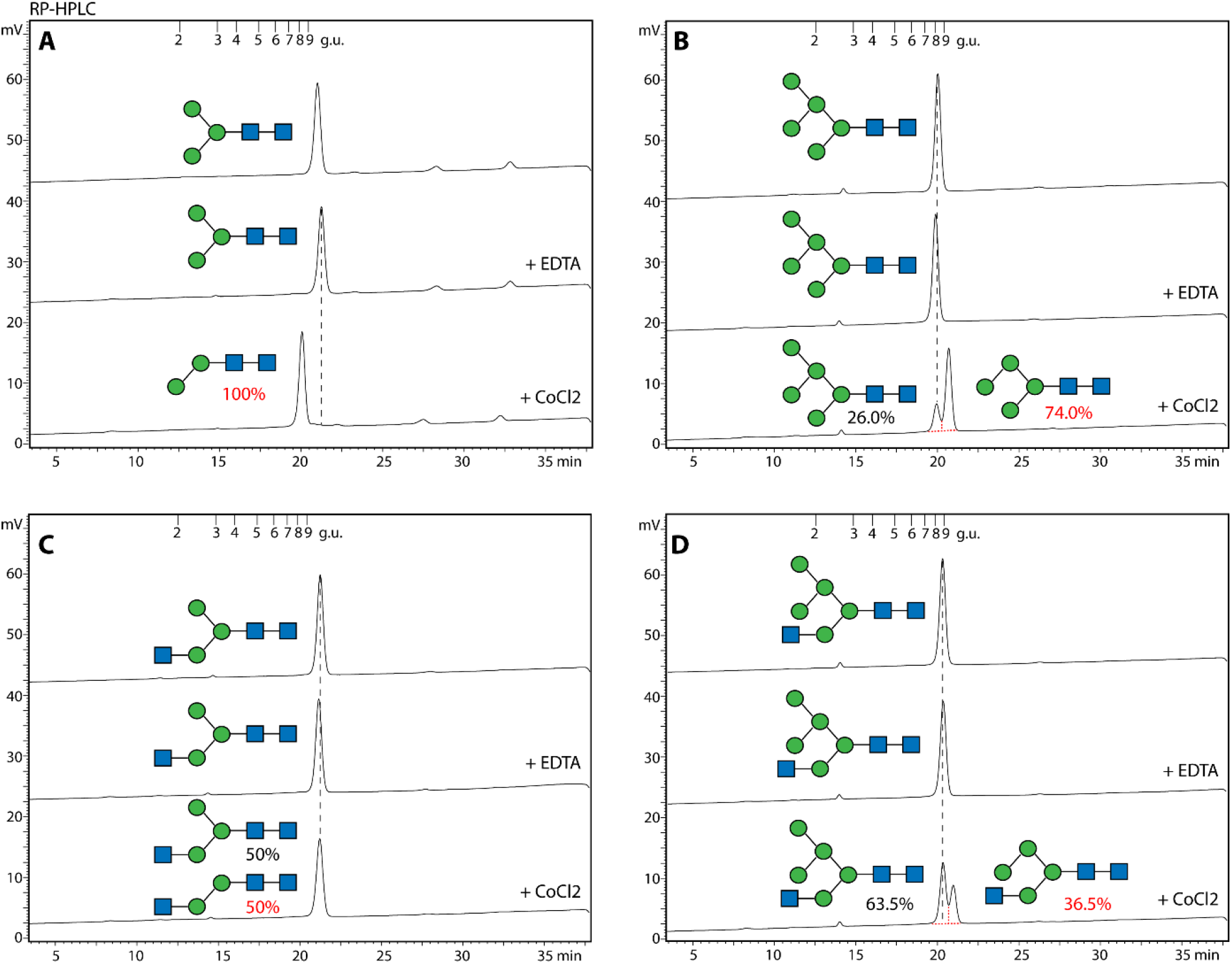
RP-HPLC quantification of AMAN-3 products. Equal amount of AEAB-labelled glycans (10 pmol each) were incubated with AMAN-3 in the presence of either EDTA or CoCl_2_. Post 16-hour incubation, all samples were heat-inactivated and analysed by HPLC on the same day. Peak areas were integrated and manually collected fractions were subject to MALDI-TOF MS analysis. Losses of one α1,6-linked mannose from Man_3_GlcNAc_2_ (**A**), Man_5_GlcNAc_2_ (**B**) and Man_5_GlcNAc_3_ (**D**) resulted minor shifts in retention times on the chromatograms. A partial conversion of Man_3_GlcNAc_3_ (**C**) to Man_2_GlcNAc_3_ was confirmed by MS data (**Supplementary Figure 5**).

### Glycan remodelling using Caenorhabditis glycoenzymes

The N-glycan biosynthesis in *C. elegans* is a series of very complicated reactions occurring in the ER and the Golgi, resulting in a large variety of structures on properly folded glycoproteins [32]. Although not yet fully understood, some of the biosynthetic pathways could be deduced from N-glycomics data obtained from wild type and knockout strains as well as from *in vitro* enzymatic data [33], whereby removal of the α1,6-mannose to result in FUT-6 substrates was considered a missing link. For the first time, we employed recombinant glycosyltransferases (GLY-13, GLY-20, FUT-1, FUT-6 and FUT-8) and glycosidases (AMAN-2, AMAN-3 and HEX-2), all recombinant forms of *C. elegans* enzymes, to mimic some of the biosynthetic reactions occurring in the Golgi by remodelling fluorescein containing glycoconjugates (***Figure 8***).

**Figure 8.**
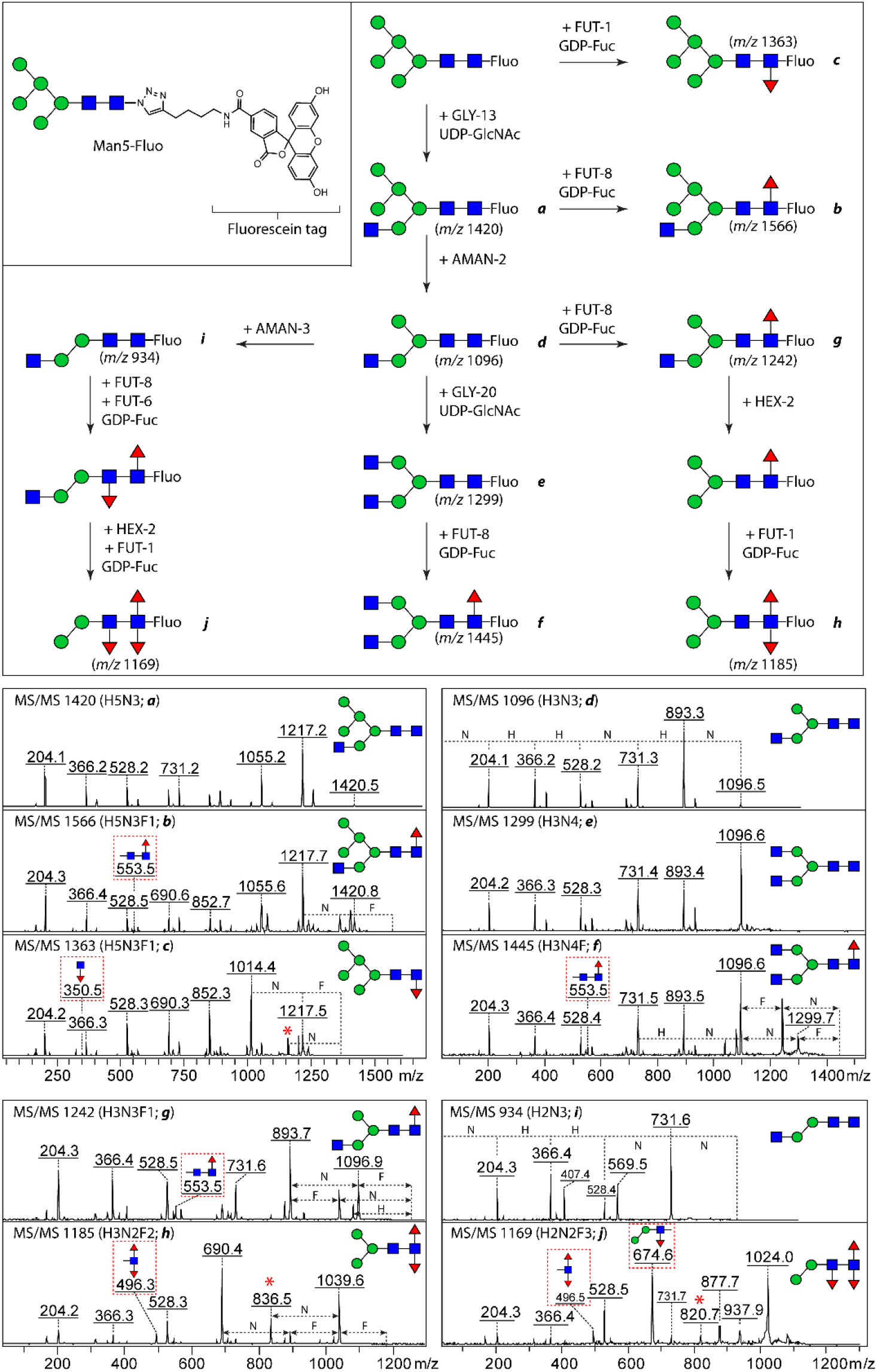
Remodelling of fluorescein-conjugated N-glycans and MALDI-TOF MS/MS spectra of products. The remodelling started with a Man_5_GlcNAc_2_ structure; five *C. elegans* glycosyltransferases as well as three glycosidases, recombinantly expressed in *Pichia pastoris*, were sequentially employed to modify their substrates; afterwards, aliquots of the reaction mixtures were analysed by mass spectrometry. A range of hybrid, complex biantennary and fucosylated paucimannosidic N-glycan structures were successfully synthesised. Structures of enzymatic products (**a** to **j**, left panel) were confirmed based on the fragmentation patterns of their parent ions ([M+H-linker]^+^), as well as the knowledge of the substrate specificities of the employed enzymes. Key fragments indicating where fucose is attached were highlighted in dashed boxes in red; whereas fragment ions, indicative of the “re-arrangement” of fucose residues, were marked with red asterisks (*e*.*g*., *m/z* 836.5).

Unlike PA-labelled and AEAB-labelled glycans, fluorescein-conjugated glycans have a tendency for in source fragmentation in MS mode; nevertheless, a portion of the intact compounds remained visible on the MS spectra. This is exemplified by analysing the intact Man5-Fluo glycan (***Supplementary Figure 1***). The MS/MS spectra of [M+H-linker]^+^ ions with a mass difference of 520 Da (loss of the linker due to in-source degradation) are, however, more informative than the spectra of the intact compounds detected as [M+Na]^+^. Therefore, [M+H-linker]^+^ ions were selected for structural assignment in MS/MS experiments and these *m/z* values are those mentioned in the following paragraphs.

Firstly, an aliquot of Man5-Fluo (5 nmol) was incubated in the presence of the donor substrate UDP-GlcNAc with GLY-13, one of three known β1,2-*N*-acetylglucosaminyltransferase I (GnT I), to yield [Manα1,6(Manα1,3)Manα1,6](GlcNAc β1,2Manα1,3)Manβ1,4GlcNAcβ1,4GlcNAc-Fluo (Man5Gn-Fluo). Full conversion was observed after overnight incubation as judged by the ion of *m/z* 1420, increased by 203 from *m/z* 1217 (***Figure 8a***); in addition, a portion of the GnT I product was fully fucosylated by a α1,6-fucosyltransferase (FUT-8), which resulted in the formation of a compound with *m/z* 1566 (corresponding to ***Figure 8b***). The core fucosylation has been confirmed by the presence of a daughter ion HexNAc_2_Fuc_1_ of *m/z* 553.5, a diagnostic ion used for other compounds carrying 6-linked core fucose residue. Under the same conditions, a small aliquot of Man5-Fluo was fully fucosylated by FUT-1 to [Manα1,6(Manα1,3)Manα1,6](Manα1,3)Man β1,4GlcNAcβ1,4(Fucα1,3)GlcNAc-Fluo and in this case its daughter ion HexNAc_1_Fuc_1_ (*m/z* 350.5, ***Figure 8c***) was more pronounced in the MS/MS spectrum. However, *C. elegans* FUT-8 cannot modify Man5-Fluo with a 6-linked core fucose *in vitro*, even though more enzyme was added and longer incubation was carried out, suggesting that this enzyme strictly requires the presence of GlcNAc on the lower arm of the reducing end. Interestingly, the observation of N-glycans with core α1,6-fucose (minor portion) in a triple GlcNAc-TI knockout indicated *in vivo* activity of FUT-8 on Man5-carrying glycoproteins [12]. This enzymatic characteristic is identical to the human homologous enzyme FUT-8 published recently [34].

Following modification by GLY-13, Man5Gn-Fluo was processed by a α-mannosidase AMAN-2 without previous purification; a new compound with *m/z* 1096 was observed after 2-hour incubation, indicative of a full conversion to Manα1,6(GlcNAcβ1,2Manα1,3)Manβ1,4GlcNAcβ1,4GlcNAc-Fluo (also known as MGn; ***Figure 8d***). Subsequently, HPLC purified MGn-Fluo was split for further remodelling into two types of N-glycans: biantennary glycans and fucosylated glycans. The former were achieved by sequential incubation with GLY-20 and FUT-8; the two products were verified by MS/MS (*m/z* 1299 and 1455, ***Figure 8e*** and ***f***). A difucosylated glycan was created by serial incubation of MGn-Fluo first with FUT-8 to introduce 6-linked fucose and then with a mixture of HEX-2 and FUT-1 to introduce a 3-linked fucose (***Figure 8g*** and ***h***). The final product carries Manα1,6(Manα1,3)Manβ1,4Glc-NAcβ1,4(Fucα1,6)(Fucα1,3)GlcNAc-Fluo (also called MMF^3^F^6^), which is a typical difucosylated N-glycan occurring in both nematodes and insects [7, 35]; this structure was concluded due to a range of daughter ions including *m/z* 496.3 Y-ion and its corresponding B-ion of *m/z* 690.2, suggesting a difucosylation on the inner most non-reducing GlcNAc residue.

In addition, by removing the ‘upper arm’ from MGn-Fluo with AMAN-3, a FUT-6 acceptable substrate was formed (***Figure 8i***). After sequential modification with core fucosyltransferases and HEX-2, removing the non-reducing GlcNAc residue, a trifucosylated glycan Manα1,3Manβ1,4(Fucα1,3)Glc-NAcβ1,4(Fucα1,6)(Fucα1,3)GlcNAc-Fluo was achieved as judged by the presence of fragments at *m/z* 496.5 and 674.6 (***Figure 8j***).

These results provided additional enzymatic data to support the sequential modifications of N-glycans by Golgi resident-glycoenzymes towards the formation of hybrid-, biantennary- and fucosylated paucimannosidic N-glycans.

### Loss of glycosylation enzymes delays development, reduces animal size and impairs food-dependent behaviours

The phenotypic characterization of *C. elegans* glyco-mutants used in this study offer an opportunity to further define the biological role of N-glycan modifying enzymes *in vivo*. All homozygous strains were viable under laboratory conditions and naked eye observation with a regular stereo microscope did not show obvious phenotypes. We thus turned to quantitative assessments of worm growth speed, size, posture, and locomotion. First, we measured the developmental speed by comparing the time from hatching to first egg-laying in wild type (N2), *bre-1, aman-3*, as well as *hex-2;hex-3* double mutants and *hex-2;hex-3;aman-3* triple mutants (***Figure 9A*** and ***B***). We found that *bre-1, aman-3* and *hex-2;hex-3* developed significantly slower than wild type (***Figure 9A***). Interestingly, the *hex-2;hex-3;aman-3* triple mutant growth was similar to that of N2 and significantly faster than that of either *aman-3* or *hex-2;hex-3*, suggesting that these mutations genetically interact to modulate developmental speed.

**Figure 9.**
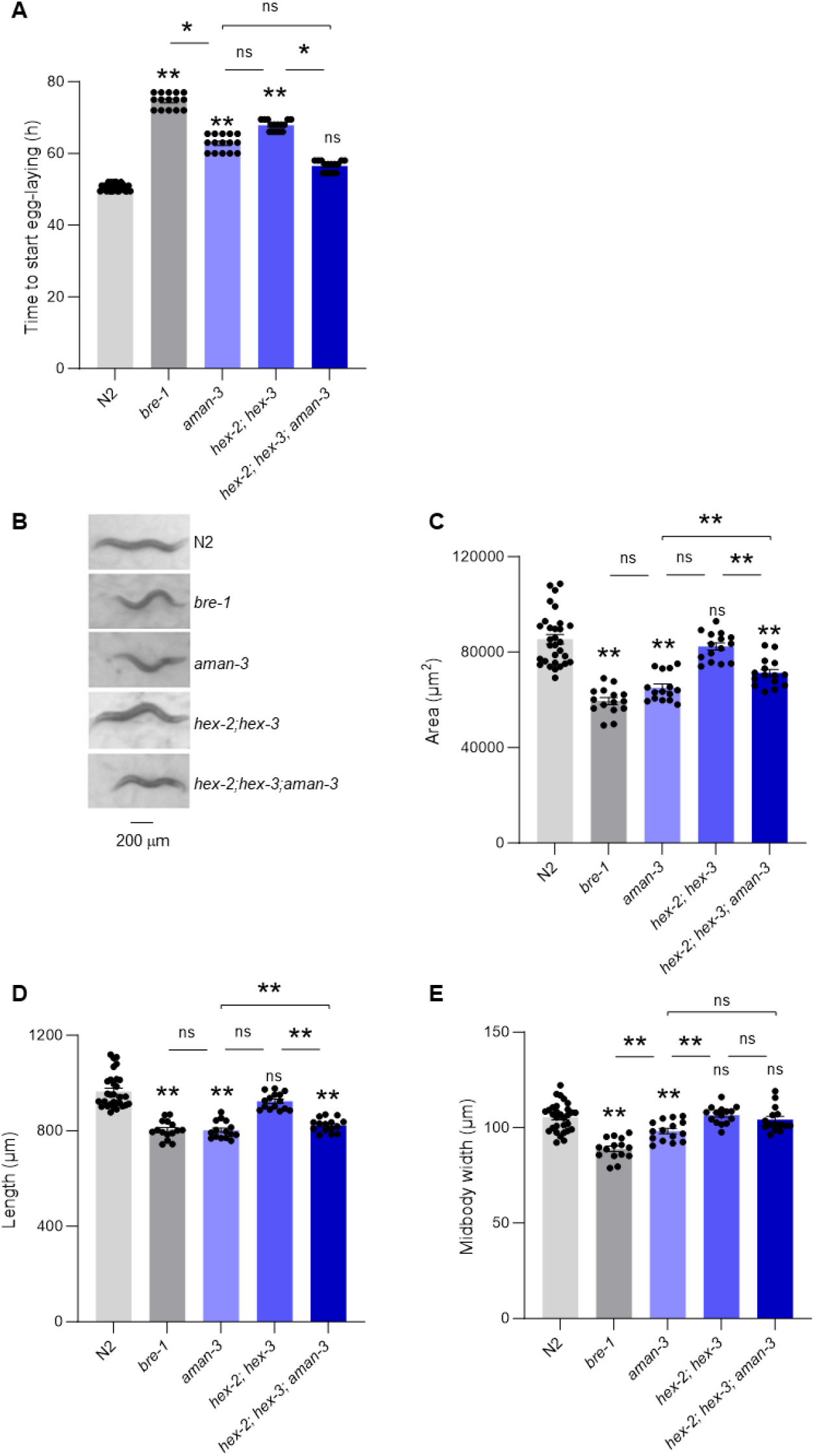
Growth and morphological defects in glyco-mutants. Comparison of developmental time (**A**) and adult animal size (**B**-**E**) between wild type (N2) and indicated glyco-mutants grown at 25°C. Average (bars) +/-s.e.m. (error bars) of *n*= 15 assays per genotype, each assay scoring at least 20 animals (**A, C**-**E**). Representative picture of adult worms illustrating size differences (**B**). **, *p*<.01; *, *p*<.05, ns, not significant by Dunn (**A**) or Bonferroni (**C**-**E**) post-hoc tests. Signs just above the glyco-mutant bars indicate significance levels versus N2.

Second, we quantified the size of adult animals (***Figure 9B-E***). Whereas the size of *hex-2;hex-3* double mutants was similar to that of wild type, it was significantly reduced in *bre-1, aman-3* and *hex-2;hex-3;aman-3* triple mutants. Both animal length and midbody width were reduced in *bre-1* and *aman-3*, whereas only length was reduced in *hex-2;hex-3;aman-3* triple mutants (***Figure 9D*** and ***E***).

Third, we used high-content computer-assisted behavioural analysis tools to reveal postural and locomotion differences across strains in crawling animals under fed and starved conditions. On food, wild type animals are in a dwelling state [36, 37], characterized by low locomotion speed, frequent pausing and foraging head movement (side-to-side nose swipes). All four mutant strains displayed altered dwelling behaviour, with up-regulated foraging movements and down-regulated pausing frequency (***Figure 11A-C***). As in previous studies [37], food-deprivation caused wild type worms to shift to a food-search behaviour characterized by increased speed (***Figure 11D***) and more frequent reorientation events (called omega turns, ***Figure 11E***), hence producing specific dispersal trajectories (***Figure 10A***), favouring longer-range exploration (***Figure 10B***). This behavioural state is also associated with a distinct posture, including increased midbody and tail bending (***Figure 11F-G***). We found that, upon food deprivation, the *bre-1* mutants and *aman-3* mutants displayed reduced speed, omega turn frequency, midbody and tail bending as compared to wild type (***Figure 11D-G***). Examples of typical postures with reduced curvature are presented in ***Supplementary Figure S6***. In contrast, *hex-2;hex-3* double mutants had an opposite phenotype with increased speed and omega turn frequency (***Figure 11D-E***). The triple *hex-2;hex-3;aman-3* mutant displayed even further increase in speed and omega-turn frequency, with values significantly increased in comparison to all the other genotypes. These specific behavioural differences across genotypes were associated with different trajectories and dispersal efficacy, that is likely relevant for food-search behaviour (***Figure 10***).

**Figure 10.**
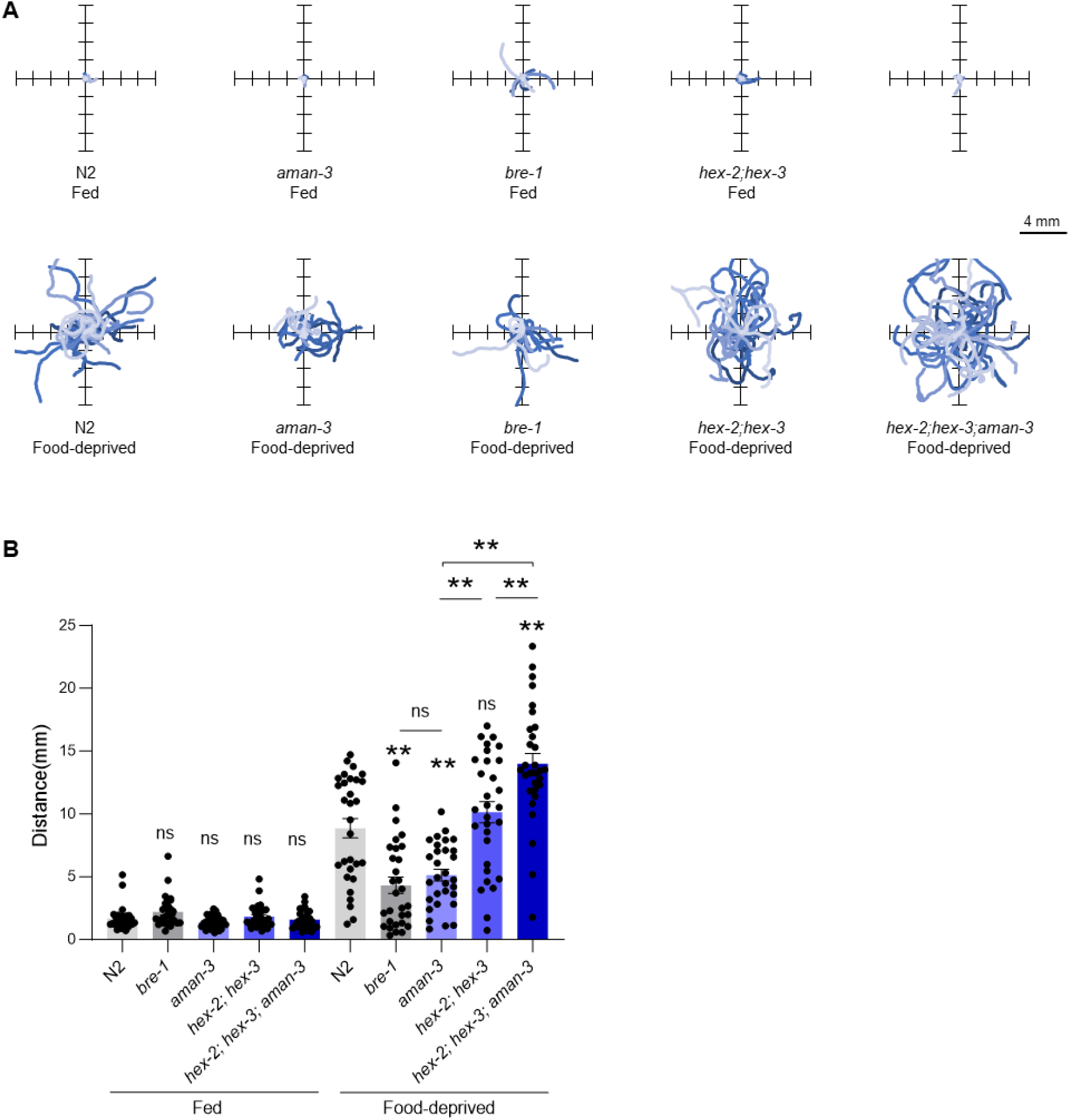
Alteration of food-search trajectories in glyco-mutants. One-minute worm trajectories of well-fed worms on food (**A**, top panels) and 3h food-deprived worms off-food (**A**, bottom panels). Thirty trajectories per condition plotted from a single starting (0,0) coordinate and illustrating the worm dispersal. Average ± s.e.m. and individual data points for the covered distance (**B**, corresponding to the path length). *n*=30 animals. *, *p*<.05 and **, *p*<.01 by Bonferroni posthoc tests. Signs just above the glyco-mutant bars indicate significance levels versus N2.

**Figure 11.**
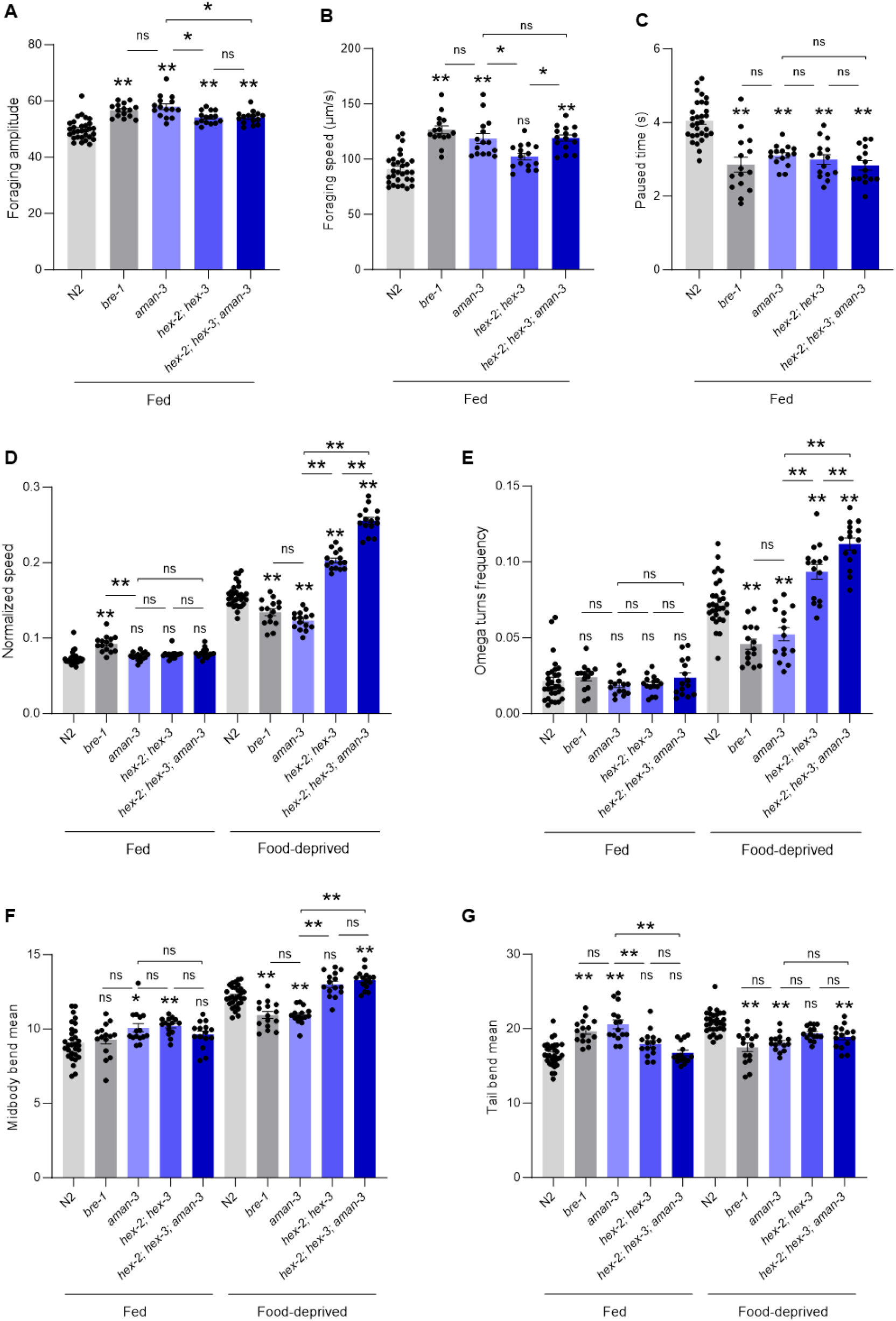
Postural and locomotion behavior alterations in glyco-mutants. Comparison of postural and locomotion parameters in adult animals between wild type (N2) and indicated glyco-mutants grown at 25°C and assessed on food (Fed, **A**-**G**) or assessed off-food 3h after food deprivation (Food-deprived, **D**-**G**). Average (bars) +/-s.e.m. (error bars) of *n*= 15 assays per genotype, each assay scoring at least 20 animals. **, *p*<.01; *, *p*<.05, ns, not significant by Bonferroni (**A**-**G**) post-hoc tests. Signs just above the glyco-mutant bars indicate significance levels versus N2.

Collectively, our phenotypic data in glyco-mutants highlights a role for *aman-3, hex-2/3* and *bre-1* in controlling worm developmental speed, animal size, dwelling behaviour on food and food-search behaviour in response to food-deprivation. Furthermore, the various phenotype-specific genetic interaction types between *aman-3* and *hex-2;hex-3* mutations suggest that the complex, non-additive impact that the losses of AMAN-3 and the two hexosaminidases have on N-glycan structures translates into pleiotropic phenotypic impact.

## Discussion

Probably all multicellular eukaryotes have multiple members of the so-called GH38 α-mannosidase family. At least one is lyososomal or vacuolar and involved in glycoconjugate degradation, whereas one or more have roles in N-glycan remodelling in the Golgi apparatus [38]. In a previous study, the *C. elegans* AMAN-3 was identified as a homologue of the Golgi α-mannosidase II (AMAN-2) and the non-purified recombinant enzyme showed to react with *p*NP-α-mannoside, an artificial substrate [23]. Here, we confirm and expand on these previous observations to substantiate the notion that AMAN-3 is a Golgi-resident α1,6-mannosidase that uses N-glycans as substrates, representing a sought-after missing link in the N-glycan biosynthetic pathway of *C. elegans* and a number of related nematode species, and filling an important gap in our understanding of the complex enzymatic orchestration into play.

AMAN-3 fits the characteristics of typical Golgi-resident mannosidases, which possess N-terminal transmembrane domains, are active at a slightly acidic pH and require metal ions as co-factors [23]. Our GFP-fusion confocal data in this study are also in line with a Golgi localisation of AMAN-3. Interestingly, AMAN-3 activity depends on the presence of Co (II) divalent metal ion. This feature is very similar to a *Spodoptera frugiperda* Golgi α-mannosidase, the SfMANIII, as well as the *Drosophila* Man-IIb (**Table 1**). Both insect and nematode enzymes can be inhibited by swainsonine, a common GH38 α-mannosidase inhibitor [21, 22]. Other Co (II)-dependent mannosidases include cytosolic or lysosomal enzymes with roles in degradation, such as mammalian MAN2C1 and MAN2B2 [39, 40].

**Table 1.**
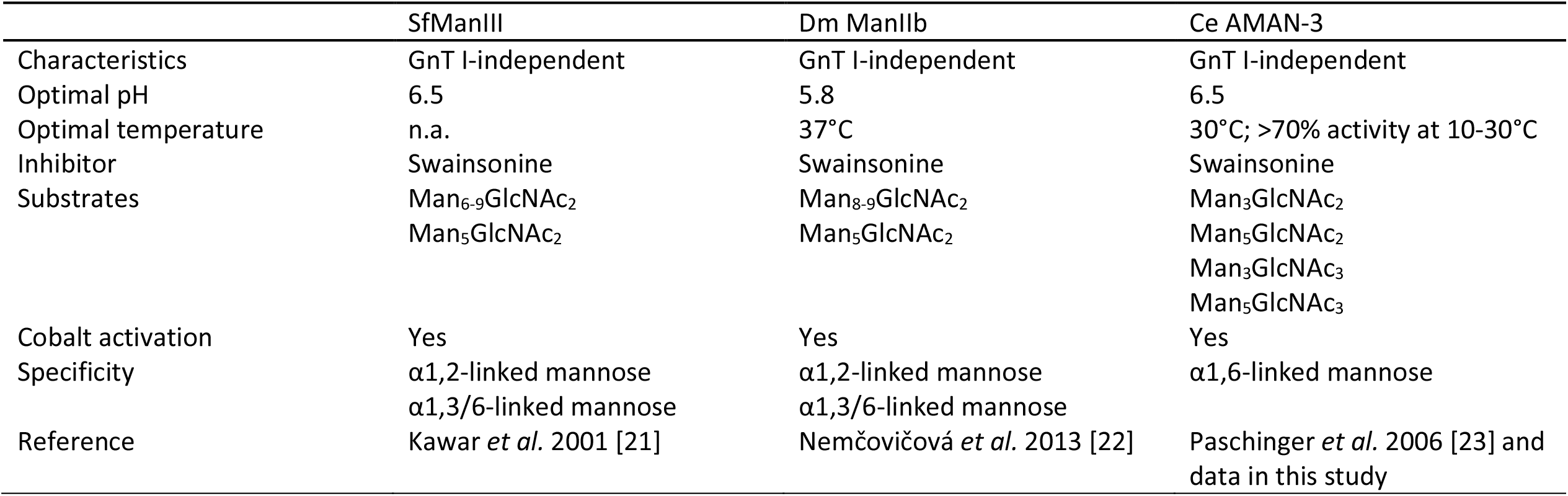
A comparison of insect and nematode Golgi α-mannosidase III.

In this study, we used reverse genetics and comparative glycomics to obtain a very detailed view of the impact of *aman-3* loss, and combined these approaches with a comprehensive *in vitro* reconstitution system of glycan remodelling able to reveal AMAN-3 action in the context of a complex substrate pool. Our glycomics data provided direct evidence that the *aman-3* gene is essential to the biosynthesis of tetra-fucosylated N-glycans occurring in the N2 wild type worms. Knocking out *aman-3* in *hex-2;hex-3* double mutant resulted in an even more dramatic shift in the N-glycome, as judged by the absence of fucose modification on the distal GlcNAc residue. One signature glycan structure in *hex-2;hex-3* with a composition of Man_2_GlcNAc_3_Fuc_2_Gal_2_ is completely abolished (***Figure 12***). Collectively, our data indicated that AMAN-3 plays a key role in removing a ‘block’ to modification by FUT-6 of the distal (second) core GlcNAc residue. Thus, it can be surmised that orthologues of AMAN-3 work in concert with orthologues of FUT-6 in species such as *H. contortus, O. dentatum* and *A. suum*, but that these are absent from species such as *D. immitis* or *T. suis* which lack trifucosylated core chitobiose motifs [20, 41, 9]. The biological role of these structures in the context of parasite-host interactions remains unclear. However, it is possible that mammalian hosts recognise these structures of parasites, as they significantly differ from mammalian glycans. Indeed, the question as to how the natural processing of parasite antigens affects their immunogenicity as well as their ability to act as protective epitopes is still open. Studies on *H. contortus* H11 antigens have indicated that recombinant forms do not induce protective immunity, even if expressed in *C. elegans* [42]; however, the ability to better reproduce natural *H. contortus* glycosylation in recombinant expression systems by employing AMAN-3 and FUT-6 may bring us closer to understanding the interplay between nematode glycosylation and host immune systems.

**Figure 12.**
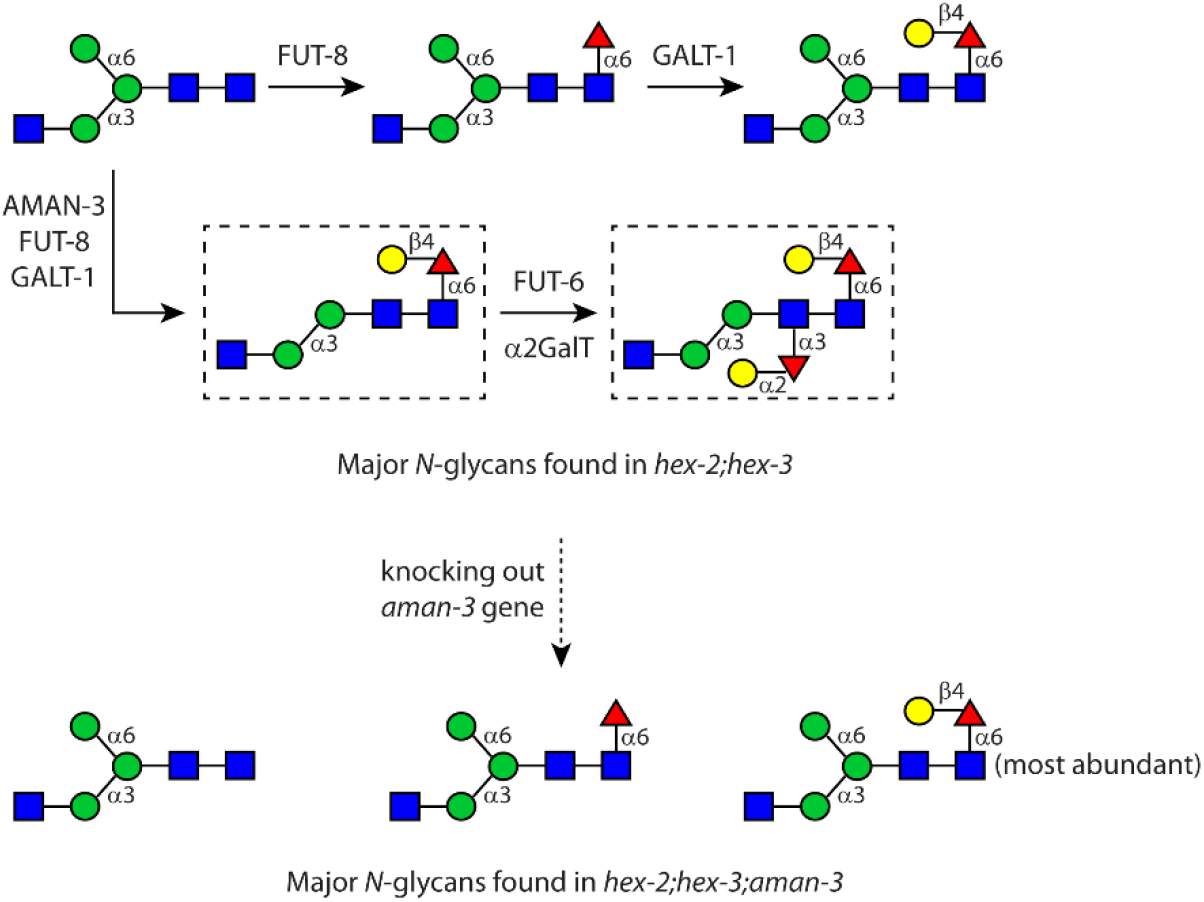
A summary of major N-glycan structures found in the double (*hex-2;hex-3*) and in the triple (*hex-2;hex-3;aman-3*) mutants. Enzymes that participate relevant biosynthetic pathways are annotated. Glycans without the ‘upper arm’ (shown in dished boxes), abundant in *hex-2;hex-3*, are abolished after knocking out the *aman-3* gene.

Beyond its likely relevance for parasite-host interaction, our study provides direct insight on the more general biological importance of AMAN-3, BRE-1 and HEX-2/3 enzymes in *C. elegans*, including their roles in growth and behaviour modulation. Interestingly, our data show that the different mutations produce sometime similar and sometimes opposite impact on the different phenotypic parameters. For instance, *bre-1* and *hex-2/-3* mutations both reduce growth speed, but their respective impact on animal speed and omega turn during food search was opposite. Likewise, the epistatic analysis between *aman-3* and *hex-2;hex-3* showed different synthetic effects, with positive epistasis effects for food-search behaviour parameters, but a negative epistasis effect for development speed. These complex phenotypic patterns are not very surprising in the light of the complex glycome alterations that these mutations produce, which are not ‘simply additive’. Indeed, each mutant produces a unique pattern of N-glycans, missing some wild-type structures and harbouring some abnormal structures (***Figure 3*** and ***4***). Behavioural differences observed in worm glyco-mutants demonstrated a close connection between protein glycosylation and developmental defects as exemplified by studies on *aman-2* and *cogc* mutants [43, 44], which is also well known in human congenital disorders of glycosylation (CDG) [45]. Our approach combining glycome data with high-content phenotypic characterization paves the road for future studies with additional *C. elegans* glyco-mutants in order to get a more comprehensive and systematic understanding of the relationships between glycome changes and their phenotypic impact in this powerful genetic model.

### Experimental procedures

*Preparation and cultivation of C. elegans strains* – The N2 wild type *C. elegans* and HY496 mutant (*bre-1* deficiency) were obtained from the Caenorhabditis Genetics Centre, University of Minnesota, MN and an *aman-3(tm5400)* single mutant, carrying a 401 bp deletion and 2 bp insertion in the *aman-3* gene, was obtained from the National Bioresource Project for the Experimental Animal Nematode *C. elegans*, Tokyo Women’s Medical University, Japan. A *hex-2;hex-3* double mutant [16] was previously prepared by crossing the single *hex-2(tm2530)* mutant and *hex-3(tm2725)* mutant, which were obtained through the National Bioresource Project; this double mutant was used to prepare a *hex-2;hex-3;aman-3* triple mutant (cop1842) using the CRISPR/Cas9 approach by Knudra Transgenics (NemaMetrix, Eugene, OR). Using two *aman-3*-targeting sgRNAs to guide Cas9, a 5145-bp fragment was deleted in the genome of the *hex-2;hex-3* mutant and replaced with a 18-bp insertion containing a 3-frame stop sequence (5’-TAAATAAATAAACTCGAG-3’). Screening was carried out by genotyping PCR using primer pairs A (5’-AGCCTCAATTTCGTCTACACCAC-3’ and 5’-GGGCAAATCGGAGTCATCTGAA-3’, 652 bp as the wild type) and B (5’-AGCCTCAATTTCGTCTACACCAC-3’ and 5’-CCTGTTTTGCACCCAGTTTGATG-3’, 724 bp as *aman-3* knockout, ***Supplementary Figure 2***) and the deletion of *aman-3* gene was confirmed by DNA sequencing. Moreover, an EGFP (enhanced green fluorescent protein)-encoding DNA fragment was fused to the *aman-3* gene of N2 worms using CRISPR/Cas9 technology. This knock-in strain (*aman-3::egfp*, strain name PHX5021; allele *syb5021*) was custom-made by SunyBiotech.

All worm strains were cultured under standard conditions at 20 °C. For a large-scale preparation, worms were grown in S-complete medium supplied with *E. coli* OP50 in shake flasks and were purified by sucrose density centrifugation; after intensive washing in saline solution, worms were stored at -80 °C prior to analysis.

*Mannosidase activity assays using worm lysate* – N2, *aman-3* and *hex-2;hex-3* worms were harvested from NGM agar plates and repeatedly washed in saline solution (0.9% NaCl) to remove OP50. Worms were transformed in a tissue grinder and homogenised in 300 µL of lysis buffer containing 20 mM MES buffer, pH 6.9, 0.5% Triton X-100 and 0.01% protease inhibitor cocktail (Sigma-Aldrich); post centrifugation to remove cell debris (14,000 g for 4 min at 4°C) clear supernatants of the worm lysates were kept for enzymatic assays. Reaction mixtures were prepared by mixing 1 µl of PA-MM substrate (2D-HPLC purified) and 0.8 µl of worm supernatant with 25 mM ammonium acetate buffer (pH 6.5) supplied with or without 6 mM cobalt (II) chloride (CoCl_2_) or 0.6 µM Swainsonine (Sigma-Aldrich). Reactions were incubated at room temperature prior to HPLC analysis.

#### Confocal microscopy

Worms at mixed developmental stages were harvested and incubated in M9 buffer supplied with 5 μM BODIPY™ TR Ceramide complexed to bovine serum albumin (Invitrogen™) at room temperature (1 hour). Prior to immobilsation on poly-L-lysine– coated glass slides, worms were thoroughly washed in M9 and paralysed in 20 mM levamisole hydrochloride. Images were recorded using a Leica TCS SP5 laser scanning confocal microscope (Wetzlar) as described previously [27].

#### Molecular cloning

The *C. elegans aman-2* CDS was subcloned from a pPICZαC construct [23] to a pPICαHisFLAG vector using the Gibson Assembly (GA) approach. GA primers (5’-CGACGATGACAAGCTGCAGGCATCGATGAAAGATGTTTGTGG-3’ and 5’-GCTGGCGGCCGCCCGCGGTTAAAATGATACAAGAATACTG-3’) were used to amplify a truncated *aman-2* gene (encoding aa 126-1145) by two-step PCR using a Q5® HIFI DNA polymerase (New England Biolabs); DNA fragment was ligated to an empty pPICαHisFLAG vector using NEBuilder® HiFi DNA Assembly Cloning Kit (New England Biolabs) and the reaction mixture was transformed to NEB5α competent cells. Followed by PCR screening, constructs of positive clones were DNA sequenced (LGC genomics). The *aman-2* construct was linearised by *Pme I*, purified and transformed by electroporation to the GS115 strain of *Pichia pastoris* for protein expression. (Note that despite previous success with an untagged AMAN-3 [23], an N-terminally tagged form was not expressed; thus, insect cell-based expression was employed as described below.)

To express AMAN-3 in insect cells, the truncated *aman-3* gene (encoding aa 35-1046) was PCR amplified (5’-GCAGCCATCAAAGATTAGGACAGCA-3’ and 5’-CGTCGACGTAGGTCATCGATAAAGAATCAA-3’) and ligated in-frame to an engineered pACEBac1 vector, which contains DNA fragments encoding a melittin signal sequence, a His-FLAG tag and a thrombin cleavage site on the N-terminus of AMAN-3. This construct is incorporated into the baculovirus genome following the instruction provided with the MultiBac™ cloning kit (Geneva Biotech).

#### Recombinant Protein Expression and Purification

Recombinant forms of *C. elegans* AMAN-2 (with or without a His-tag), AMAN-3 (non-tagged), GLY-13 (non-tagged), GLY-20 (non-tagged) as well as His-tagged HEX-2, FUT-1, FUT-6 and FUT-8 were expressed in *Pichia pastoris* for 72-96 hours at 16°C following the manufacturer’s manual. Methanol was added to the culture daily to maintain an induction concentration at 1%. Harvested culture supernatants were concentrated and buffer-exchanged by ultrafiltration using 30 kDa MWCO centrifugal devices (Sartorius, Germany). Hand-packed nickel affinity columns (Qiagen, Hilden, Germany) were used to obtain purified recombinant proteins that carry a N-terminal poly-histidine tag (AMAN-2, HEX-2 and FUTs).

His-tagged AMAN-3 was expressed in Sf9 insect cells. Briefly, cells were maintained in HyClone™ SFM4Insect medium (GE Healthcare) supplied with 3% of fetal bovine serum (Gibco™) at 27°C. Post transfection of Sf9 cells with a Bacmid DNA (1 µg) using 10 µl of FuGene (Promega), recombinant baculoviruses carrying *aman-3* gene were harvested (V_0_) and used to infect Sf9 cells in a 68 ml suspension culture (80 rpm, 27°C). 3 days post infection, the recombinant AMAN-3 was purified from the cell culture supernatant by His-tag purification. The protein sequence was verified by LC-MS/MS at the MS core facility of the University of Veterinary Medicine Vienna.

#### Mannosidase activity assays using recombinant enzymes

To assess the biochemical property of AMAN-3, including metal ion dependency, temperature and pH optima, a colorimetric assay using *p-* nitrophenyl-α-mannopyranoside as a substrate (pNP-Man, dissolved in dimethyl sulfoxide) and Co(II) was employed as previously described [23]. Briefly, His-Tag-purified AMAN-3 was incubated with 5 mM pNP-Man in quadruplicate in a 96-well plate under various conditions using McIlvaine buffers. Post terminating the reactions with 250 μl of 0.4M glycine-NaOH, pH 10.4, optical densities at 405 nm (OD405) were measured using a Tecan Infinite M200 micro-plate reader.

To generate N-glycan substrates for testing recombinant mannosidases, a Man_8_GlcNAc_2_ structure conjugated with 2-amino-N-(2-aminoethyl)benzamide (*AEAB*, purchased from NatGlycan LLC) was remodelled. 10 nmol of this compound was first digested with α1,2-mannosidase (yielding Man_5_GlcNAc_2_) and then processed by GLY-13 in the presence of donor substrate UDP-GlcNAc and Mn (II) (yielding Man_5_GlcNAc_3_). Man_3_GlcNAc_3_ and Man_3_GlcNAc_2_ structure were prepared by digesting Man_5_GlcNAc_3_ with AMAN-2 alone or in combination with HEX-2. All resulted compounds were HPLC purified and quantified based on the integrated areas of corresponding eluent peaks.

#### Release of N-glycans and PA-labelling C. elegans

strains at mixed-stages (4-6 g in wet weight) were boiled in water to denature endogenous proteases and glycosidases; worms were homogenised in a tissue grinder prior to a 2-hour proteolysis at 70 °C in a round-bottom glass flask using thermolysin (Promega, 1 mg per 1 g wet-weight worm) in 50 mM ammonium bicarbonate buffer (pH 8.5) supplied with 0.5 mM CaCl_2_ [25]. Post centrifugation to remove insoluble cell debrides, glycopeptides were enriched by cation exchange chromatography (Dowex 50W×8; elution with 0.5 M ammonium acetate, pH 6.0) and desalted by gel filtration (Sephadex G25, 0.5% acetic acid as solvent). N-linked glycan were released using a recombinant *Oryza sativa* PNGase A (New England Biolabs) in 50 mM ammonium acetate buffer (pH 5.0) overnight at 37°C. Native glycans were separated from residual peptides by cation exchange chromatography (Dowex 50W×8); glycans collected in the filtrate fraction (as no longer bund to the column) were further purified using hand-packed C18 cartridges and nPGC cartridges prior to analysis. Native glycans were labelled with 2-aminopyridine (PA) to induce a fluorescent tag at the reducing ends and the excess reagent was removed by gel filtration (Sephadex G15, 0.5% acetic acid as solvent) as previously described [25].

#### Glycan remodelling

In total, eight active enzymes (AMAN-2, AMAN-3, GLY-13, GLY-20, HEX-2, FUT-1, FUT-6 and FUT-8) were employed for the remodelling experiments. All recombinant enzymes were expressed in-house as described above. Except for His-tag enzymes that were purified, crude enzymes post concentration and buffered-exchange against a storage buffer (25 mM Tris-HLC, 150 mM NaCl, pH 7.0) were used. A fluorescein-conjugated Man_5_GlcNAc_2_ structure (Man5-Fluo) was chemically synthesised and verified by NMR (detailed in **Supplementary Information**).

In brief, 5 nanomoles of Man5-Fluo were used as the initial acceptor substrate, sequentially modified by glycosyltransferases and glycosidases according to the experimental design (***Figure 8***). For assays using glycosyltransferases, mixtures containing 80 mM MES buffer (pH 6.5), 20 mM MnCl2, substrate, 2 mM nucleotide sugar and relevant enzyme were prepared, whereas for assays using glycosidases, mixtures containing 160 mM MES buffer, substrate and relevant enzyme were prepared. Incubation was carried out overnight at 37°C to ensure the full conversion, except for AMAN-2 which was incubated for 2 hours to reduce a by-product. Reaction mixtures were directly analysed by MALDI TOF MS and a few reaction products were HPLC purified prior to MS analyses to obtain improved MS and MS/MS data.

#### HPLC methods

A Shimadzu Nexera UPLC system equipped with a RF 20AXS fluorescence detector was used to analyse and fractionate different glycoconjugates. For N-glycomics studies, PA-labelled worm glycans were fractionated over a Ascentis® RP-amide column (2.7 µm, 15 cm × 4.6 mm attached to a 5 cm guard column) on HPLC. A gradient of 30% (v/v) methanol (buffer B) in 0.1 M ammonium acetate, pH 4.0 (buffer A) was applied at a flow rate of 0.8 ml/min as follows: 1% buffer B per minute over 35 minutes (excitation/emission: 320 nm/400 nm) [17]. In addition, separation of PA-glycans were also achieved using a Hypersil ODS column (Agilent) with the same buffers and detector settings, but a flow rate of 1.5 ml/min.

A HyperClone reversed phase column (5µ ODS C18, 250×4 mm; Phenomenex) was used for the separation and quantification of AEAB-labelled N-glycans and the purification of fluorescein-conjugated glycans. AEAB-glycans: 0.1 M ammonium acetate, pH 4.0 as buffer A and 30 % (v/v) methanol as buffer B, and an optimised gradient as follows: 0–10 min, 0-20 % B; 10–35 min, 20–50 % B; 35–35.5 min, 50 % B; 35.5–36 min, 0% B; 36–40 min, back to starting conditions. The flow rate was set at 1.5 ml/min with a maximal pressure at 375 bars and the fluorescence detector setting was Ex 330 nm and Em 420 nm. Fluorescein-glycans: using 0.1% formic acid as buffer A and a mixture of 99.9% acetonitrile and 0.1% formic acid as buffer B; detector setting was Ex 490nm and Em 500nm. All HPLC fractions were manually collected, dried and further examined by MALDI TOF MS.

#### MALDI-TOF MS and MS/MS

lyophilised glycan samples, either in native or derivatised forms (PA, AEAB and fluorescein conjugates), were dissolved in HPLC grade water and subject to MALDI-TOF MS and MS/MS analyses on an Autoflex Speed instrument (1000 Hz Smartbeam-II laser, Bruker Daltonics) using 6-aza-2-thiothymine as a matrix [46]. Calibration of the instrument was routinely performed using the Bruker Peptide Calibration Standard II to cover the MS range between 700 and 3200 Da. The detector voltage was normally set at 1977 V for MS and 2133 V for MS/MS; typically, 3000 shots from different regions of the sample spots were summed. Automatic measurements were conducted on HPLC-fractionated glycan samples (N2, tm5400 and cop1842) using the AutoXecute feature of the control software (Bruker flexControl 3.4). To obtain MS/MS spectra, parent ions were fragmented by laser-induced dissociation (LID) without applying a collision gas (precursor ion selector was generally ±0.6%). For PA and AEAB-labelled glycans, [M+H]^+^ ions were favoured for LID fragmentation, whereas for fluorescein conjugates [M+H-linker]^+^ ions, as ‘in-source degradation’ products of the intact compounds, were fragmented because their MS/MS spectra presented the most informative fragments for structural annotation.

#### Growth, morphological and behavioural analyses

Animal synchronization was made by treating gravid adults with standard hypochlorite-based procedure. Developmental speed was assessed by measuring the time between hatching and the onset of egg laying in animals maintained at 25°C on regular NGM plates seeded with *E. coli* OP50. For each genotype, the growth time prior to behavioral assays was adjusted in order to test animals just after they started to lay their first eggs. Animal size, postural and locomotion parameters were obtained from videos recorded and analyzed as in Thapliyal et al. [37] using the Tierpsy Tracker v1.4 [47]. Fifteen assays per condition, each tracking a population of at least 20 animals, were recorded and the average value of selected postural and locomotion parameters extracted for each assay. Individual data points (dots) overlaid in the different bar graphs each represent the value derived from one assay. The ‘food-deprived’ condition was acquired 3 hours after food-deprivation. Locomotion speed was normalized to the body length in order to compensate for variable animal sizes across genotypes. D’Agostino & Pearson test (p < 0.01) was used to test normality of distributions. Comparisons giving significant effects (p < 0.05) with ANOVAs were followed by Bonferroni posthoc tests. Dunn’s test was used as non-parametric test whenever the normality assumption criterion was not fulfilled. All tests were two-tailed.

## Supporting information

Nematode AMAN-3_Suppl Table 1

## Supplementary Information

## Further information regarding the glycomic analyses

### Definition of the level of the glycan structural analysis

To compare compositional differences of the overall N-glycomes of *C. elegans* N2 wild type and knockout stains, native N-glycans enzymatically released from worm materials were profiled using MALDI-TOF mass spectrometry. To gain structural details, the N-glycomes of N2, tm5400 and *hex-2;hex-3;aman-3* were subject to HPLC fractionation and MALDI-TOF MS/MS experiments.

### Search parameters and acceptance criteria

a. **Peak lists:** As stated in the methods section: typically 1000 shots were summed for MALDI-TOF MS and 5000 for MS/MS. Spectra were processed with the manufacturer’s software (Bruker Flexanalysis 3.3.80) using the SNAP algorithm with a signal/noise threshold of 6 for MS (unsmoothed) and 3 for MS/MS (four-times smoothed).
b. **Search engine, database and fixed modifications:** All glycan data were manually interpreted and no search engine or database was employed; the fixed modification is the 2-aminopyridine label at the reducing end (GlcNAc1-PA fragments of *m/z* 300).
c. **Exclusion of known contaminants and threshold:** All glycan data were manually interpreted; only peaks with an MS/MS consistent with a pyridylaminated chitobiose core were included – the ‘threshold’ for inclusion was an interpretable MS/MS spectrum (at least in terms of composition).
d. **Enzyme specificity:** A description of the PNGase Ar release method is given in the methods section; the enzyme should remove N-glycans from glycopeptides regardless of the presence of core α1,3-fucose on the reducing-terminal GlcNAc.
e. **Isobaric/isomeric assignments:** For isomeric species, differences in RP-HPLC elution and MS/MS were used for the assignment (as described in the text).

### Glycan or glycoconjugate identification

a. **Precursor charge and mass/charge (*m/z*):** All glycans detected were singly-charged. For the positive mode, the *m/z* values are for protonated forms. Depending on the glycan amount or preparation, the relative amounts of the H^+^, Na^+^ and K^+^ adducts varied. Maximally two decimal places used for the *m/z* annotations consistent with the accuracy of MALDI-TOF MS; in the figures and due to space limitations, only one decimal place is presented. Previous data indicate an average +0.03 Da (+ 22 ppm) deviation between the measured and the calculated *m/z* values on the instrument used.
b. **MALDI-TOF MS settings (positive mode):** Ion Source 1 and 2 were 19.00 and 16.75 kV; Lens, 9.00 kV; Reflector 1 and 2, 21.05 and 9.65 kV; Pulsed Ion Extraction, 160 ns; Matrix Suppression typically up to 700 Da; Detector Gain, typically 2163 V.
c. **MALDI-TOF MS/MS settings (positive mode):** Ion Source 1 and 2 were 6.00 and 5.35 kV; Lens, 2.90 kV; Reflector 1 and 2, 27.00 and 11.75 kV; Lift 1 and 2, 19.00 and 4.00 kV; Pulsed Ion Extraction, 140 ns; Detector Gain, typically 2260 V when fragmenting; Laser Power Boost typically 50%; not in CID mode; PCIS typically 0.65%.
d. **All assignments:** For the glycans present in each pool, see the RP-HPLC chromatograms annotated with structures shown according to the Standard Nomenclature for Glycans.
e. **Modifications observed:** Listed are the *m/z* values for glycans carrying a reducing terminal pyridylamine group as judged by presence of an *m/z* 300 GlcNAc1-PA fragment. As the glycans are otherwise chemically unmodified, *Δm/z* of 146, 160, 162, 176, 165 and 203 correspond to fucose, methyl-fucose, hexose, methyl-hexose, phosphorylcholine or *N-*acetylhexosamine (positive ion mode). There was no indication for the presence of sulphate, phosphate or sialic acid residues.
f. **Number of assigned masses:** Glycan assignments were not just based on measured mass only, but on the basis of MS/MS corroborated by elution data (*i*.*e*., glucose unit of HPLC separation).
g. **Spectra:** Representative annotated spectra (MS and MS/MS) defining structural elements are given in various figures. In total, MS and/or MS/MS data for approximately 40 defined structures are shown.
h. **Structural assignments:** As noted in the results section, the typical oligomannosidic structures are assigned based on elution time and fragmentation pattern; it is otherwise assumed that the glycans contain a di- or tri-mannosyl core consistent with typical eukaryotic N-glycan biosynthesis and that there is processing by GlcNAc-transferases (GnTI, II and V) and core α1,3/6-fucosyltransferases as in a range of multicellular organisms.

### Synthesis of fluorescein-labelled Man5 (Man5-Fluo)

a. 20.0 mg (42.2 μmol, 1 eq) of 5-carboxyfluorescein-OSu **1** (Sigma-Aldrich) and 16.9 mg (127 μmol, 3 eq) of hexinylamine hydrochloride **2** were dissolved in 844 μL of DMSO and 70.3μL (506 μmol, 12 eq) of triethylamine were added. After 24 h at ambient temperature the mixture was dried in high vaccum. The residue was dissolved in 20 mL of water/acetonitrile 1:4 containing 0.1 % of formic acid. Two portions of 10 mL were purified by solid phase extraction (Waters SepPak C18 Classic, 2×360 mg). Each portion of 10 mL was loaded on to the cartridges follwed by washing with 10 mL of water/acetonitrile 1:4 containing 0.1 % of formic acid. The amide eluted with 10 mL of water/acetonitrile 2:3 containing 0.1 % of formic acid. After lyophilization 11.3 mg of 5-carboxyfluorescin-hexinylamide **3** (24.8 μmol, 58.8 %) were obtained. C_27_H_21_NO_6_ (455.47). LC-MS (RP18, 0-20 % CH_3_CN/H_2_O + 0.1 % formic acid): M_calcd_ = 455.14; M_found_ = 456.04 (M+H)^+^.
b. 1 μL (320 nmol) of TGTA **4** (325 mM in water) and 1 μL (320 nmol) of tetrakis(acetonitrile)copper(I)-hexafluorophosphate (325 mM in DMSO) were mixed under argon and incubated for 30 min. 1.0 mg (794 nmol, 1 eq) of **Man5-N**_**3**_ and 1.08 mg (2.38 μmol, 3 eq) of alkine **3** were dissolved in 8 μL of water under argon. Subsequently, 0.5 μL of the solution containing the preformed Cu-TGTA-complex (80 nmol) were added. After keeping the reaction under argon for 24h the mixture was diluted with 1 mL of water and purified by solid phase extraction (Waters SepPak C18 Classic, 2×360 mg). The solution was loaded on to the cartridges follwed by washing with 10 mL of water. The product was eluted with water/acetonitrile 1:7 containing 0.1 % of formic acid). After lyophilization the procedure was repeated. Yield: 770 μg of **Man5-Fluo** (449 nmol, 56.5 %), C_73_H_98_N_6_O_41_ (1715.59). LC-MS (RP18, 0-20 % CH_3_CN/H_2_O + 0.1 % formic acid): M_calcd_ = 1714.58; M_found_ = 1715.27 [M+H]^+^.

Scheme of chemical synthesis of fluorescein-labelled Man5 and the 1H-NMR spectrum of compound **3**

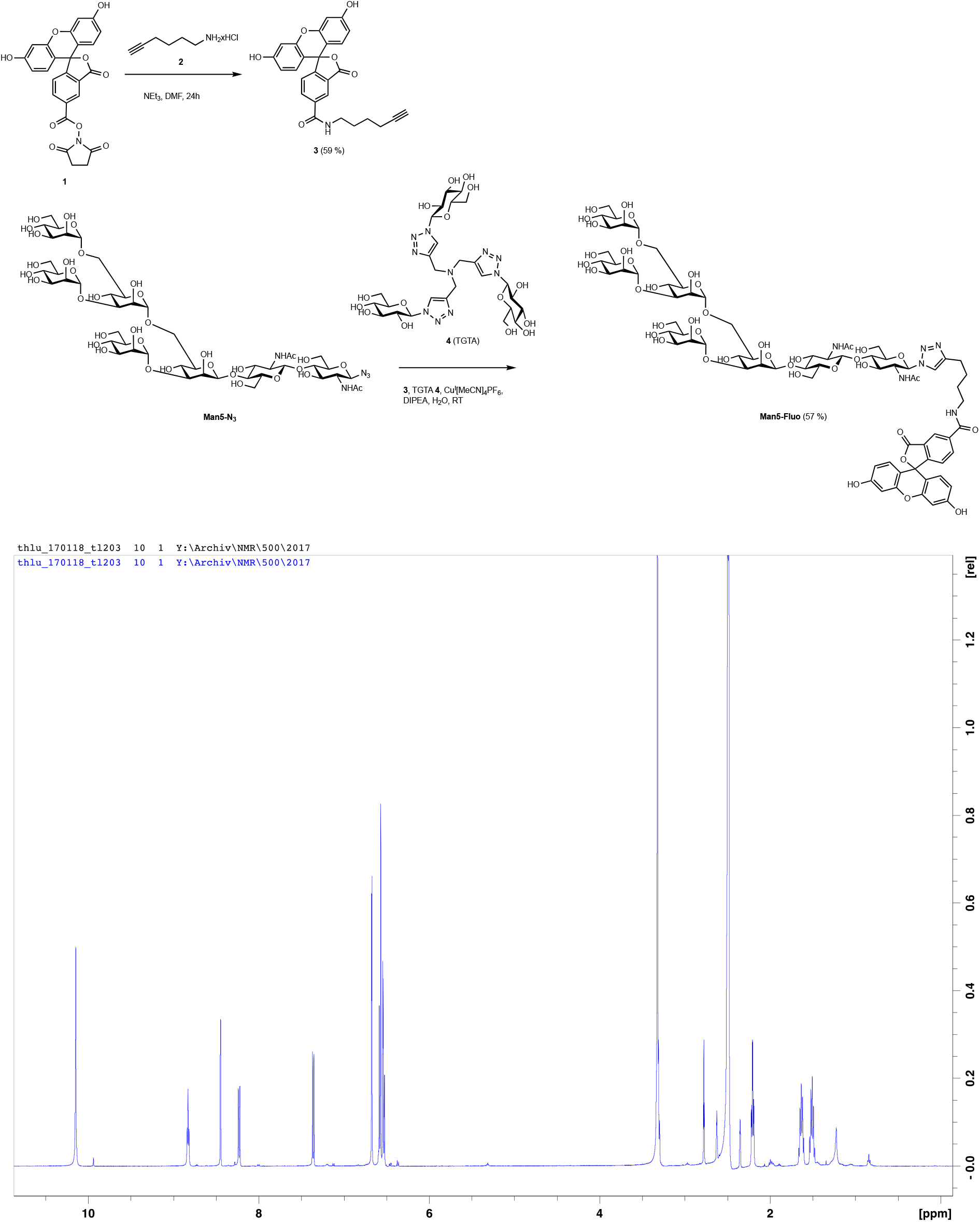

**Supplementary Figure 1.**
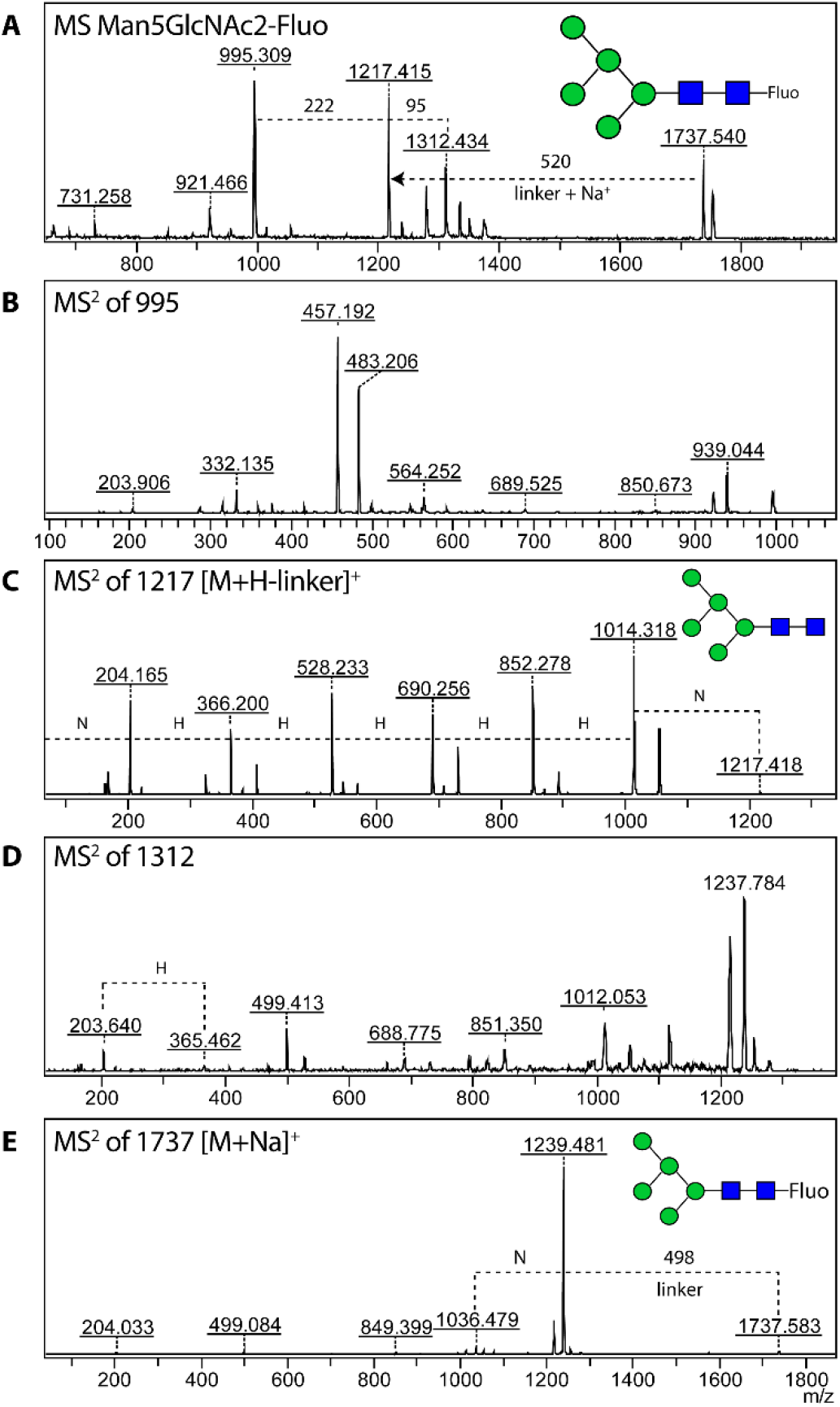
MALDI-TOF mass spectrometric properties of Man5-Fluo in the positive ion mode. MS spectrum of the intact Man5-Fluo (*m/z* 1737.5, [M+Na]^+^) displayed a series of ions (**A**), possibly due to the “laser-induced degradation”. In comparison to the other MS^2^ spectra of the same compound (**B, D and E**), the fragmentation of the *m/z* 1217.4 ion resulted in the most informative spectrum (**C**, [M+H-linker]^+^), revealing structural details of the glycan portion; whereas fragmentation of the intact sodiated compound (**E**, [M+Na]^+^) resulted in a dramatic loss of the linker (Δ = 498) as well as minor ions indicative of the composition of the glycan. This phenomenon was observed on the other fluorescein-labelled glycans in the remodelling experiment (**Figure 8**), which became the reason for choosing the [M+H-linker]^+^ ions of fluorescein-labelled compounds for MS^2^ fragmentations. N, HexNAc; H, Hexose.

**Supplementary Figure 2.**
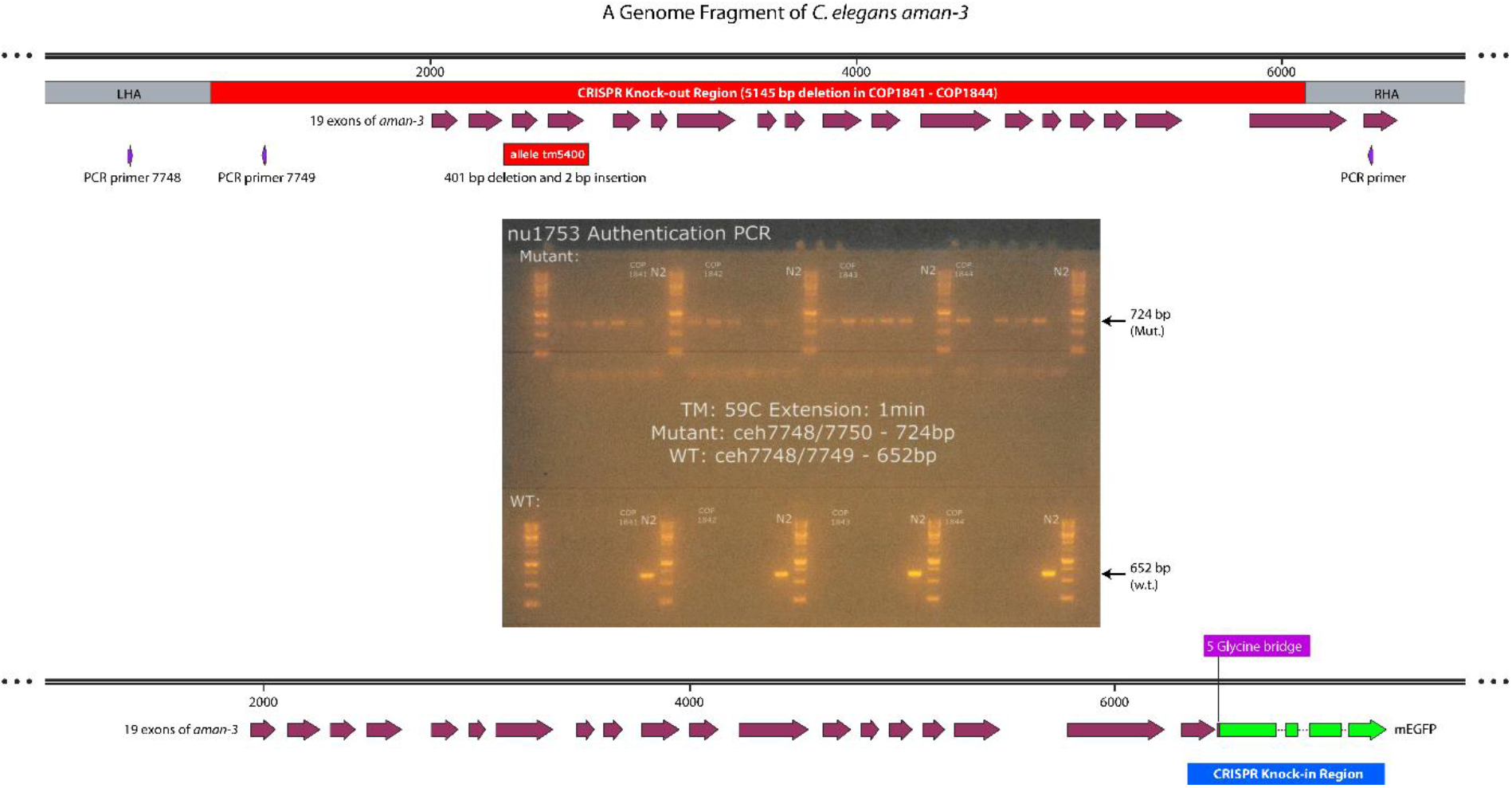
The gene structure of *C. elegans aman-3*. Deletions in a single mutant (*tm5400*) and triple knockout strains (*hex-2;hex-3;aman-3*, cop1841-cop1844) are marked in red. The CRISPR knock-in region with an insertion of mEGFP-encoding fragment (*egfp* gene being fused to the 19^th^ exon of *aman-3* via a 5×glycine bridge) is marked in blue. CRISPR mutants were verified by and DNA sequencing.

**Supplementary Figure 3.**
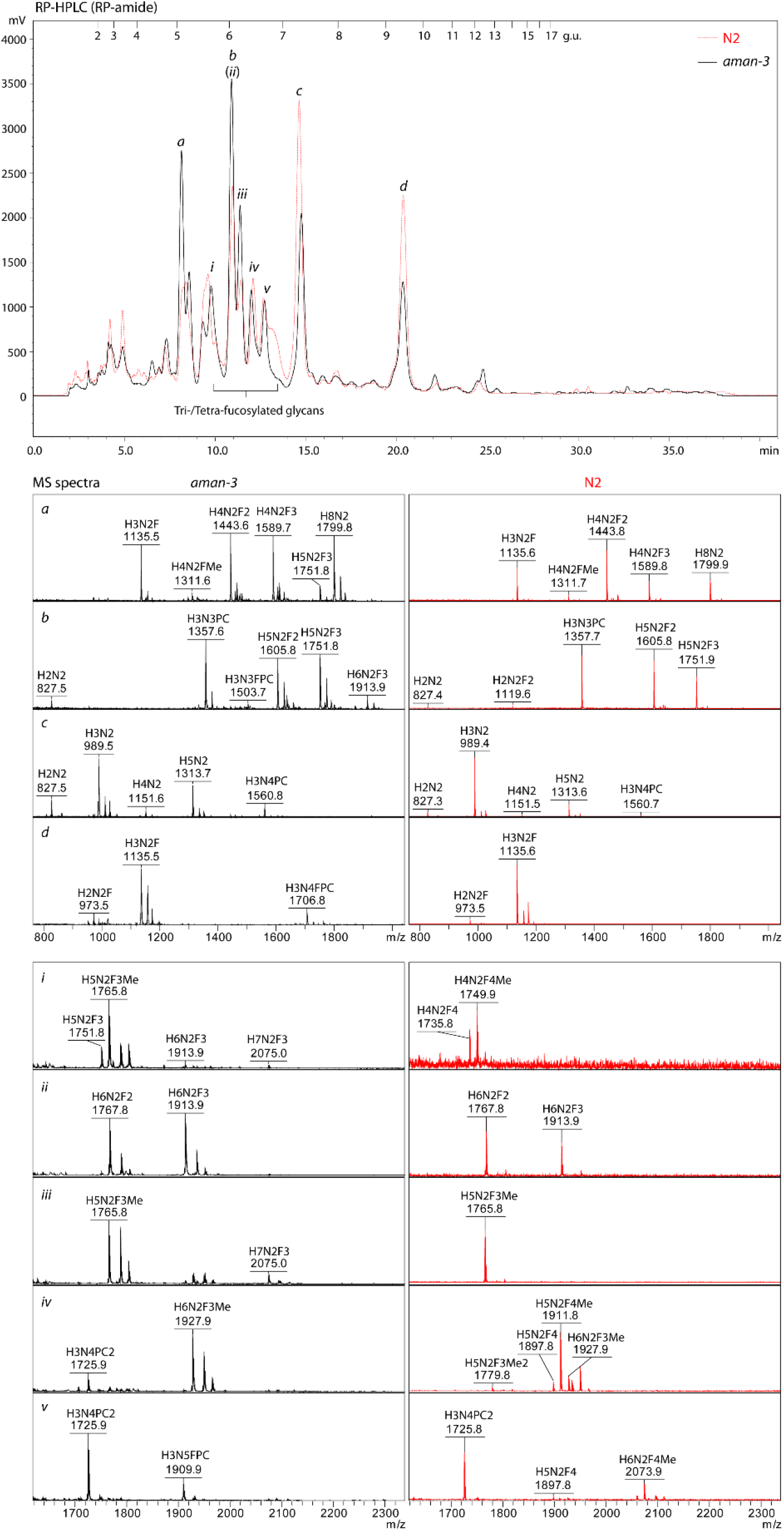
Alignment of HPLC chromatograms of N2 and tm5400 N-glycans. Aliquots of PA-labelled N-glycans, released by PNGase A, were resolved by a RP-amide reversed phase column on HPLC using dextran oligomers as a standard. N2, dashed line in red; tm5400, solid line in black. MALDI MS spectra of major HPLC fractions (*a, b, c* and *d*) and tri-/tetra-fucosylated-glycan enriched fractions (*i* to *v*; between 5.6 and 7.0 g.u.) are shown, indicative of the absence of tretrafucosylated structures in tm5400.

**Supplementary Figure 4.**
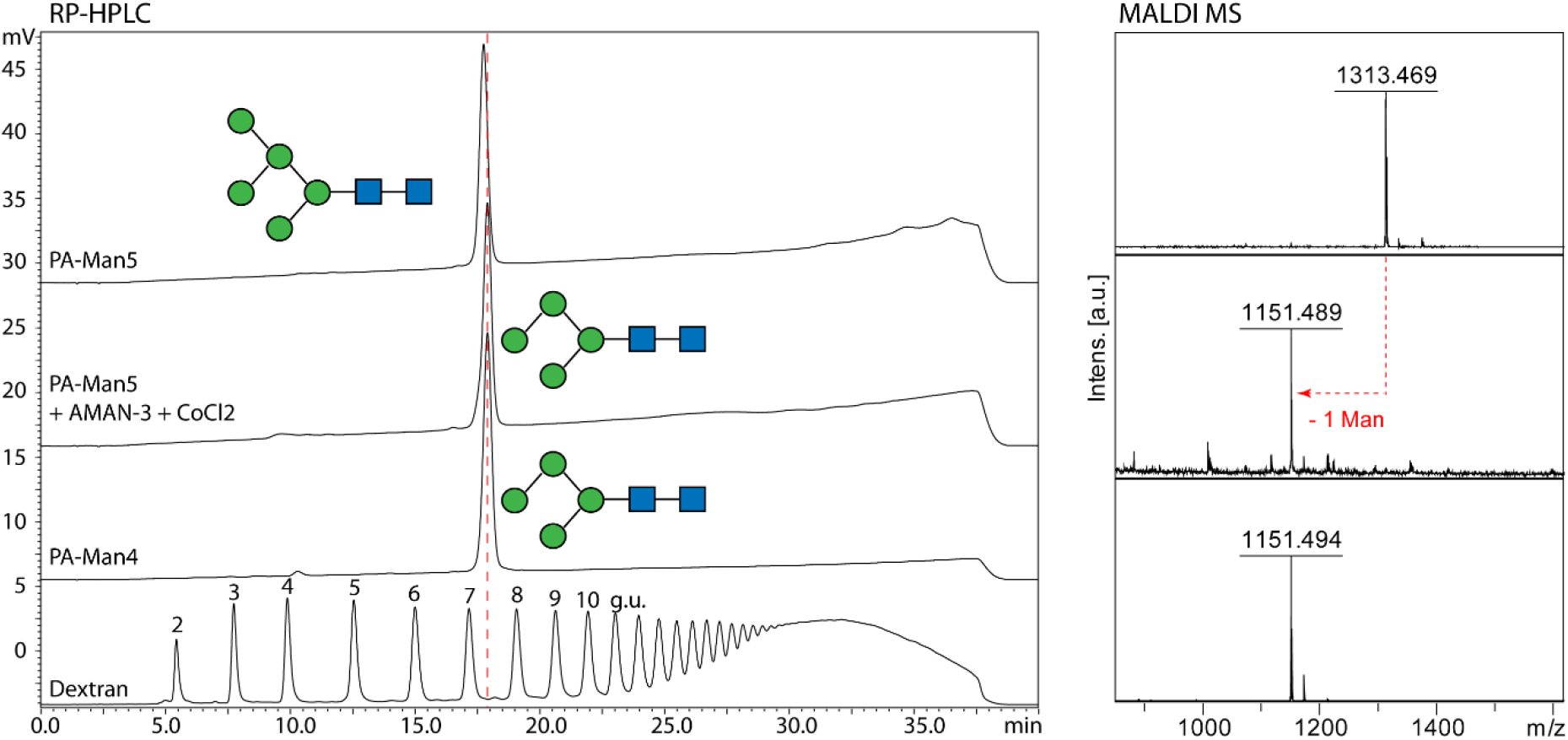
C18 RP-HPLC analysis of a PA-glycan treated with AMAN-3. An aliquot of Man_5_GlcNAc_2_ structure (PA-Man5) purified from a triple GnTI mutant (Trigly) was digested by AMAN-3 and the reactive product displayed a small shift from 17.5 minute to 18.0 minute on the HPLC chromatogram, which co-eluted with a previously characterised structure Man_4_GlcNAc_2_ (PA-Man4) of the Trigly mutant [49]. The loss of one mannose residue from PA-Man5 was confirmed by MALDI-TOF-MS.

**Supplementary Figure 5.**
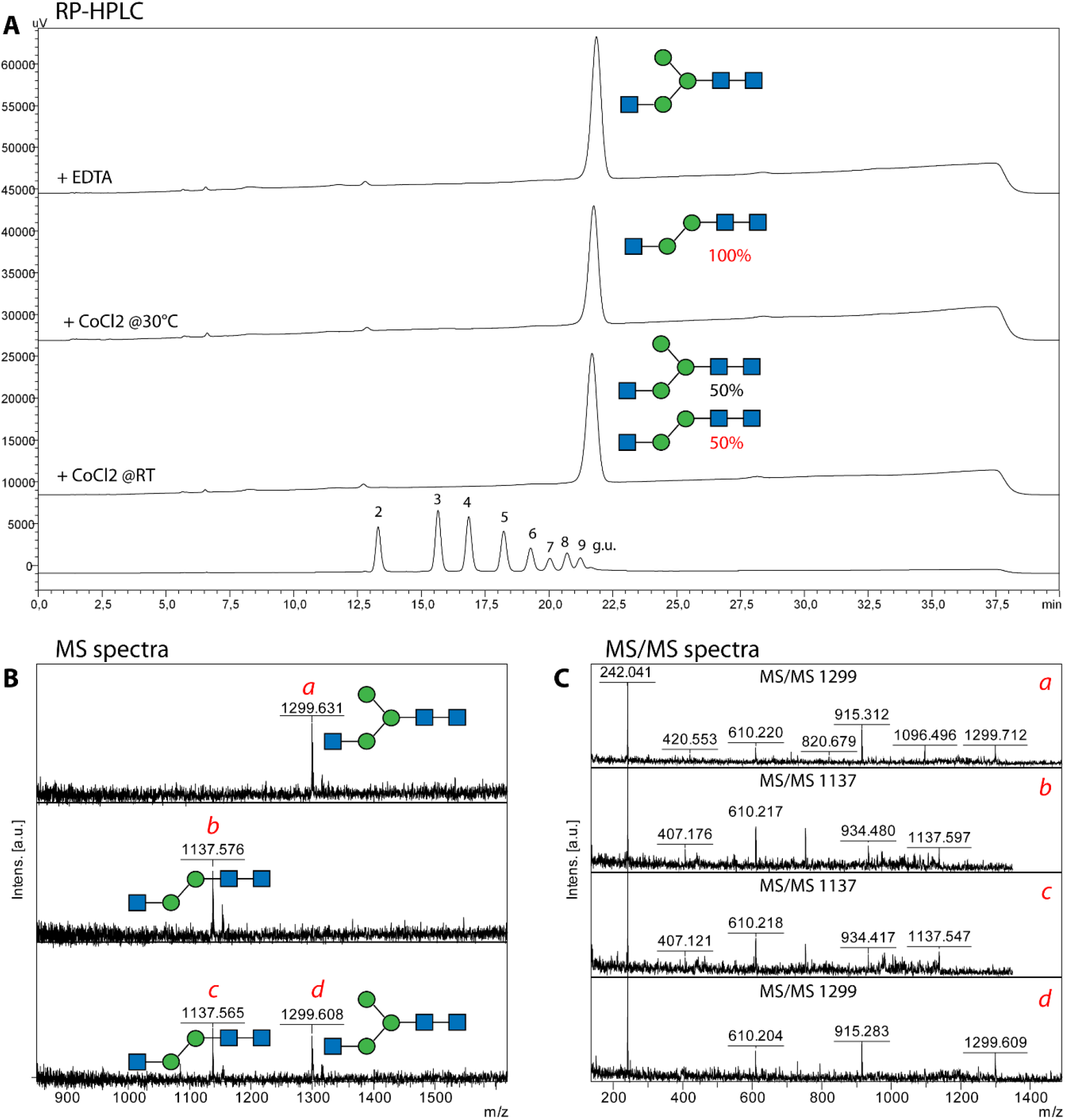
Co-elution of AEAB-labelled glycans on RP-HPLC. Man_3_GlcNAc_3_ (MGn)was incubated with AMAN-3 in the presence of CoCl2 either at 30°C or at room temperature. Post heat-inactivation, reaction mixtures were analysed on RP-HPLC (**A**) and subsequently, all eluents were examined by MALDI TOF MS/MS (**B** and **C**). Full conversion from Man_3_GlcNAc_3_ to Man_2_GlcNAc_3_ was observed in the sample incubated at 30°C whereas at room temperature, approximately 50% of MGn was digested by AMAN-3 (MS spectra *b-d* in **B**). AEAB-labelled MGn and Man_2_GlcNAc_3_ co-eluted at 21.5 minute.

**Supplementary Figure 6.**
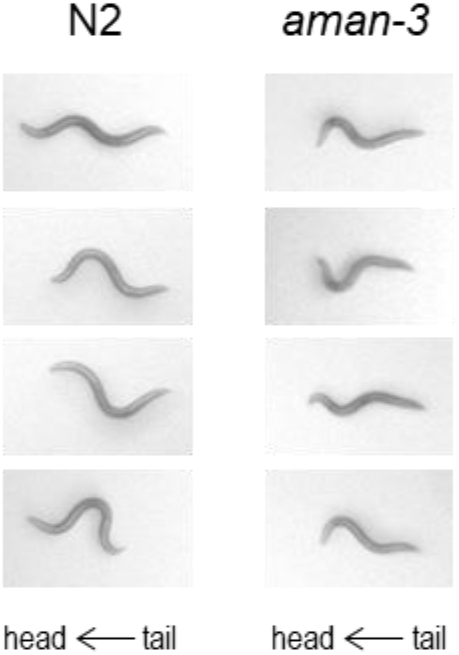
Illustration of body curvature differences between N2 and *aman-3* mutants. Snapshots from behavioral movies illustrating animal posture differences between food-deprived N2 and *aman-3* mutants. For each genotype, a single population image from a single assay was randomly selected. The presented illustrative close-up pictures were manually chosen to highlight the difference between a subset of typically N2 postures and specific examples of aman-3 mutants where midbody and tail curvature is obviously reduced.

## Data availability

Relevant MS and MS/MS data are converted to mzxml files and uploaded to GlycoPost: https://glycopost.glycosmos.org/entry/GPST000395

Preview URL: https://glycopost.glycosmos.org/preview/824990868669939fabd25c

pin code: 8263

## Acknowledgments

This work was supported by the Austrian Science Fund (FWF; grants P30021 to S.Y., P29466 to I.B.H.W and P32572 to K.P.); The Swiss National Science Foundation (grant 310030_197607 to DAG). We are thankful to the Japanese National Bioresource Project for providing *C. elegans* mutants; some strains were obtained from the CGC, which is funded by NIH Office of Research Infrastructure Programs (P40 OD010440). The authors acknowledge Dr. Zuzanna Dutkiewicz and Dr. Nicola Palmieri for their help with predicting putative glycoenzymes in nematode genomes, Licha Wortha for PCR verification of genotypes. We thank the BOKU Core Facility Mass Spectrometry for the use of the Autoflex Speed MALDI-TOF MS instrument and Dr. Karin Hummel at the VetCore Proteomics for protein ID services.

## Author contributions

S.Y., J.K., F.W., S.T., E.A., H.G., S.K., D.M., M.S.,D.P. and T.L. methodology; S.Y., S.T., D.A.G. and K.P. formal analysis; S.Y., I. B. H. W. and K.P. data curation; S.Y. and I. B. H. W. writing–original draft; S.Y., S.T., D.M. and C.U. visualization; D.A.G., S.Y. and I. B. H. W. supervision.

